# A non-canonical role of somatic CYCLIN D/CYD-1 in oogenesis and reproductive aging, dependent on the FOXO/DAF-16 activation state

**DOI:** 10.1101/2023.12.09.570929

**Authors:** Umanshi Rautela, Gautam Chandra Sarkar, Ayushi Chaudhary, Debalina Chatterjee, Mohtashim Rosh, Aneeshkumar G. Arimbasseri, Arnab Mukhopadhyay

**Affiliations:** Molecular Aging Laboratory, National Institute of Immunology, Aruna Asaf Ali Marg, New Delhi 110067, India; Molecular Genetics Laboratory, National Institute of Immunology, Aruna Asaf Ali Marg, New Delhi 110067, India; Current address: Department of Pediatrics, Washington University School of Medicine, St. Louis, MO, USA

**Keywords:** Cyclin D/CYD-1, FOXO/DAF-16, insulin signaling, pachytene arrest, germ line, somatic gonad

## Abstract

For the optimal survival of a species, an organism coordinates its reproductive decisions with the nutrient availability of its niche. Thus, nutrient-sensing pathways like insulin-IGF-1 signaling (IIS) play an important role in modulating cell division, oogenesis, and reproductive aging. Lowering of the IIS leads to the activation of the downstream FOXO transcription factor (TF) DAF-16 in *Caenorhabditis elegans* which promotes oocyte quality and delays reproductive aging. However, less is known about how the IIS axis responds to changes in cell cycle proteins, particularly in the somatic tissues. Here, we show a new aspect of the regulation of the germline by this nutrient-sensing axis. First, we show that the canonical G1-S cyclin, *cyclin D*/*cyd-1*, regulates reproductive aging from the uterine tissue of wild-type worms. Then, we show that knocking down *cyd-1* in the uterine tissue of an IIS receptor mutant arrests oogenesis at the pachytene stage of meiosis-1 in a FOXO/DAF-16-dependent manner. We find that activated FOXO/DAF-16 destroys the somatic gonad tissues like the sheath cells, and transcriptionally prevents the spermatogenesis-to-oogenesis switch to orchestrate this arrest. Deleting FOXO/DAF-16 releases the arrest and restores the somatic gonad but leads to the production of poor-quality oocytes. Together, our study reveals the unrecognized cell non-autonomous interaction of CYD-1 and FOXO/DAF-16 in reproductive aging and the regulation of oogenesis.

## Introduction

Germline is the most precious tissue of an organism that ensures perpetuation of a species. However, with aberrant metabolism brought about by aging and metabolic disorders, the reproductive tissues deteriorate leading to poor quality of oocytes, adversely affecting reproductive outcomes and progeny fitness (Jones and Lane, 2012; May-Panloup et al., 2016; Meldrum et al., 2016; Sanchez-Garrido and Tena-Sempere, 2020; Sasaki et al., 2019). To ensure optimal growth of the gametes and the fertilized oocytes, the process of oogenesis takes inputs from other reproductive and somatic niches in the body. For example, oogenesis is firmly regulated by the nutrient-sensing (Jouandin et al., 2014; Lopez et al., 2013; Tang and Han, 2017) and stress-responsive pathways (Meiselman et al., 2018; Perkins et al., 2016). In *C. elegans*, the insulin/IGF-1 signaling (IIS) pathway, a conserved neuro-endocrine signaling axis that is central to nutrient sensing and stress resilience (Evans et al., 2008; Henderson and Johnson, 2001; McColl et al., 2010; Murphy et al., 2003), regulates both somatic (Kenyon et al., 1993; Tatar et al., 2003) as well as reproductive aging (Hughes et al., 2007; Luo et al., 2010a) in a cell- autonomous as well as non-autonomous manner (Apfeld and Kenyon, 1998; Iser et al., 2007; Luo et al., 2010a; Michaelson et al., 2010; Qin and Hubbard, 2015). In mammals, IIS is crucial for the activation, and growth of the primordial oocyte follicles (Edson et al., 2009; Louhio et al., 2000; Poretsky and Kalin, 1987). The imbalance in IIS underpins pathophysiologies of multiple ovarian dysfunctions including PCOS and infertility (Azziz et al., 2016; Diamanti-Kandarakis et al., 2008; Poretsky et al., 1992; Wu et al., 2014). The IIS-PI3K-AKT axis, when activated, maintains the FOXO transcription factor in the cytoplasm through inhibitory phosphorylation (Biggs et al., 1999; Brunet et al., 1999; Paradis and Ruvkun, 1998). The overactivation of the PI3K-AKT pathway that is downstream of the IIS receptor has been shown to cause global activation of the oocyte follicles and depletes the ovarian reserve, leading to premature reproductive aging and infertility in mice (Reddy et al., 2008). On the other hand, when the signaling through the IIS pathway is low, FOXO is released from its cytoplasmic anchor and enters the nucleus to activate gene expression (Biggs et al., 1999; Brunet et al., 1999; Murphy et al., 2003; Paradis and Ruvkun, 1998). It has been noticed that FOXO activation during the early stages of oocyte development preserves the ovarian reserve and extends reproductive capacity in mice (Castrillon et al., 2003; Pelosi et al., 2013). In *C. elegans*, where the mechanisms of aging are relatively well worked out (Olsen et al., 2006; Uno and Nishida, 2016; Zhang et al., 2020) mutations in the *daf-2* (the IIS receptor ortholog) lower signaling flux through this pathway, leading to a long life and a slower reproductive aging (Hughes et al., 2007; Luo et al., 2010a; Templeman et al., 2018) The *daf-2* worms have a delayed decline in oocyte quality (Luo et al., 2010b; Templeman et al., 2018) and germline stem cell pool with age (Qin and Hubbard, 2015). Interestingly, activated DAF-16 is required in the intestine and muscle for oocyte quality maintenance (Luo et al., 2010b) while DAF-16 in the somatic gonad delays age-related germline stem cell loss (Qin and Hubbard, 2015). It is important to note that in most cases, activated FOXO is pro-longevity and it preserves reproductive fidelity.

To identify unexplored interactors of the IIS pathway, we had earlier performed a reverse genetic screen and identified a cyclin-dependent kinase CDK-12 that regulates oogenesis cell non-autonomously (Sarkar et al., 2023). To broaden our understanding of other cell cycle regulators that may have non-canonical roles in aging and if they crosstalk with the IIS pathway, we screened for cyclins that regulate germline development. Interestingly, this led to the identification of *cyclin D*/*cyd-1* (this study), a core cell-cycle protein that regulates the G1 to S phase transition (Baldin et al., 1993; Park and Krause, 1999). Beyond their involvement in cell cycle progression, cyclins and CDKs exhibit diverse biological functions (Hydbring et al., 2016; Palmer and Kaldis, 2020). Cyclin D is particularly noteworthy for its involvement in a broad spectrum of biological functions, encompassing metabolic regulation (Abella et al., 2005; Phelps and Xiong, 1998), organ development (Cenciarelli et al., 1999; Kiess et al., 1995), DNA damage repair (Jirawatnotai et al., 2011), and transcriptional control (Adnane et al., 1999; Iwatani et al., 2010; Ratineau et al., 2002).

In this study, we report two novel and unanticipated observations: 1) *cyclin D*/*cyd-1* knockdown (KD) solely in the somatic uterine tissues leads to accelerated reproductive aging and poor oocyte quality in the germline, cell non-autonomously, and 2) *cyclin D/cyd-1* KD in the uterine tissues causes FOXO/DAF-16-dependent arrest of the germline in the pachytene stage of meiosis-I specifically in the *daf-2* mutant (where DAF-16 is in an activated state). We also show an unexpected role of activated DAF-16 in inducing dramatic somatic gonad defects and downregulation of genes important for the spermatogenesis-to-oogenesis switch that may cause the germline pachytene arrest and halt oogenesis.

Together, our study highlights the non-canonical, cell non-autonomous function of *cyclin D* in reproductive aging and the unconventional role of activated DAF-16 in causing tissue damage to stall oogenesis. Considering the conserved nature of these players, it is tempting to speculate that such mechanisms may be conserved during evolution.

## Results

In a recent study, we described the crosstalk of the cyclin-dependent protein kinase, *cdk- 12* with the IIS pathway to regulate germ cell development (Sarkar et al., 2023). Interestingly, in that study, the function of the *cdk-12* was found to be independent of its canonical cyclin, *cyclin K/ccnk-1* (Blazek et al., 2011). Following up on this study, we asked if other cyclins may crosstalk with the IIS pathway to regulate germline development. For this, we knocked down (KD) the annotated cyclins in *C. elegans* in *daf-2(e1370)* (referred to as *daf-2*) and found that the RNAi knockdown of *cyclin D/cyd-1 and cyclin E/cye-1*, core cell-cycle proteins that regulate G1 to S phase transition, caused a DAF-16 dependent sterility specifically in a *daf-2* mutant (**Figure S1A**). To determine the effect of *cyd-1* and *cye-1* KD on germline development, the gonads of day-1 adults were stained with DAPI to mark the nucleus. The KD of *cye-1* drastically reduced the germ cell number in *daf-2* worms, as is expected from its known function in cell division (**Figure S1B**), whereas *cyd-1* KD demonstrated a non-canonical role in germ cell development without dramatic changes in mitosis, as described below. The loss-of-function *cyd-1* alleles are arrested during early larval development and are sterile (Park and Krause, 1999). Consequently, they could not be used for our experiments to study interaction with the IIS pathway and so, we focused our efforts on investigating the role of *cyd-1* in germ cell development and quality assurance using RNAi.

### *Cyclin D/cyd-1* maintains oocyte quality and prevents reproductive aging in wild-type worms

Since *cyd-1* KD renders *daf-2* worms sterile, we first asked how *cyd-1* may influence wild-type (WT) germline where insulin signaling may be considered normal. The *C. elegans* hermaphrodite gonad has two U-shaped tubular arms where the germ line stem cell (GSC) pool resides near the distal end and divides mitotically. On moving away from the distal end, the germ cells enter meiotic prophase. The sperm formation takes place at L4 stage (sperms are stored in the spermatheca), after which a switch from sperm-to-oocyte fate takes place. Subsequently, the germ cells are committed to developing into oocytes (Austin and Kimble, 1987; Kraemer et al., 1999; McCarter et al., 1997a). The quality of oocytes is ensured by programmed cell death of defective germ cells at the end of pachytene region. A common uterus that carries the fertilized eggs (until they are laid) connects both the gonadal arms (Kimble and Crittenden, 2007) (**Figure 1A**; only one gonad arm is shown).

**Figure 1.**
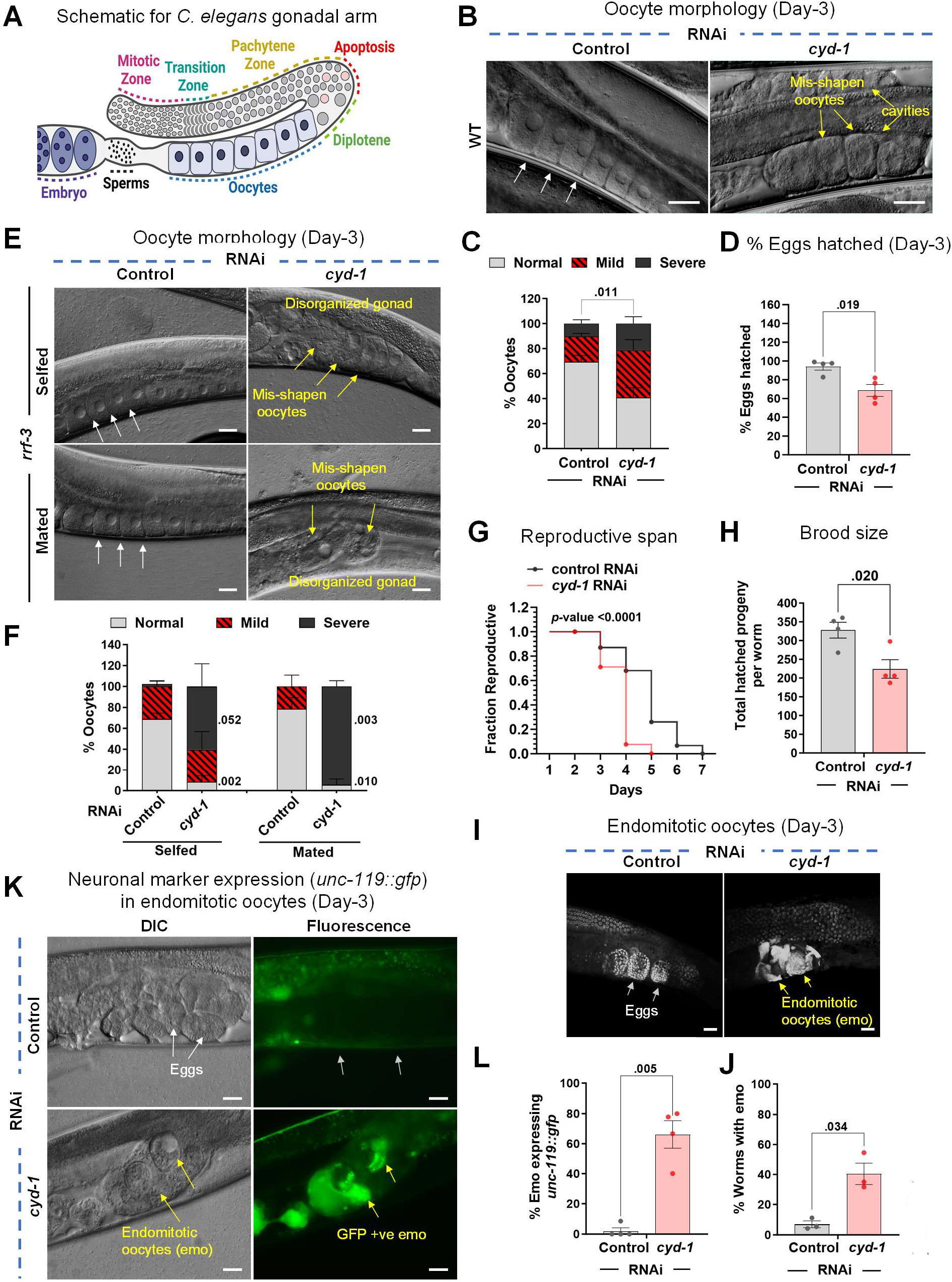
CYD-1 ensures oocyte quality and regulates reproductive aging in wild-type. **(A)** A diagrammatic representation of the right arm of *C. elegans* gonad. **(B,C)** DIC images showing oocyte morphology of WT (day-3 adult) grown on control or *cyd-1* RNAi. White arrows mark normal oocytes while yellow arrows mark poor-quality of oocytes that have cavities or are misshapen (B). Oocyte quality scores (based on morphology) (C). The quality was categorized as normal, or mild or severe based on its morphology (cavities, shape, organization) as shown in Figure S1G. Average of three biological replicates (n ≥ 25 for each replicate). Unpaired *t*-test with Welch’s correction. **(D)** Percentage of WT eggs (day-3 adult) that hatched on control or *cyd-1* RNAi. Average of four biological replicates (n ≥ 50 for each replicate). Unpaired *t*-test with Welch’s correction. **(E,F)** Oocyte morphology in unmated or mated day-3 adult worms grown on control or *cyd-1* RNAi (E). Oocyte quality scores (based on morphology as shown in Figure S1G) (F). Average of three biological repeats. Unpaired *t*-test with Welch’s correction. **(G)** The reproductive span of WT on control or *cyd-1* RNAi. **(H)** The total number of hatched progenies in WT worms grown on control or *cyd-1* RNAi. Average of four biological repeats. Unpaired *t-*test with Welch’s correction. **(I,J)** Representative DAPI-stained gonads of WT (day-3 adult) worms grown on control or *cyd-1* RNAi. White arrows mark normal eggs while yellow arrows mark endomitotic oocytes (emos) (I). Quantification of percent endomitotic oocytes (J). Average of three biological replicates (n ≥ 25 for each experiment). Unpaired *t*-test with Welch’s correction. **(K,L)** Representative fluorescent images of *unc-119::gfp* worms (day-3 adult) grown on control or *cyd-1* RNAi. White arrows mark normal eggs that do not show *gfp* expression, while yellow arrows mark endomitotic oocytes that express *gfp* (K). Quantification of *unc-119::gfp* expression positive endomitotic oocytes (L). Average of four biological replicates (n ≥ 25 for each experiment). Unpaired *t*-test with Welch’s correction. Scale bars:20 μm. Error bars are s.e.m. Experiments were performed at 20°C. Source data are provided in Table S1.

WT worms were grown from larval stage 1 (L1) onwards on control or *cyd-1* RNAi, DAPI stained at day-1 and imaged to observe the germ cell number (**Figure S1C,D**). Interestingly, we did not observe changes in the WT mitotic or transition zone germ cell number but found a significant reduction in the pachytene cell numbers (**Figure S1D**). We questioned if a decrease in the pachytene germ cell number could be due to an increase in germ cell apoptosis. Indeed, we observed an increase in apoptotic germ cell numbers marked as *ced-1::gfp*-expressing foci in the gonad [CED-1 is a transmembrane protein on the surface of sheath cells that engulfs the apoptotic germ cells, thereby marking the dying germ cells (Zhou et al., 2001)] in *cyd-1* RNAi worms (**Figure S1E,F**). Normally, as the pachytene cells transition to diplotene, most of the defective/damaged germ cells are culled in the turn region of the gonad to ensure that only healthy oocytes mature (Andux and Ellis, 2008). The increase in apoptosis upon *cyd-1* KD suggested an increase in damaged germ cells. We therefore evaluated the oocyte quality as a readout of germ cell quality. Firstly, the oocytes were examined morphologically by using differential interference contrast (DIC) microscopy (magnification of 400 X) on day-3 of adulthood. The oocytes were assigned a score (normal, mild or severely defective morphology) based on the presence of cavities, abnormal shape/size, and/or organization (**Figure S1G**). At day-3 of adulthood, WT worms grown on control RNAi showed the proper arrangement of stacked oocytes without any cavities (**Figure 1B,C**). However, upon *cyd-1* KD, the oocytes became significantly more misshapen, and the gonads were disorganized, with cavities between oocytes (**Figure 1B,C**). Further, we asked if the observed poor oocyte morphology also translates into poor quality of these oocytes. Poor quality of oocytes often leads to a reduction in the hatching efficiency of the eggs. In the case of the wild- type worms, the percentage of hatched eggs is known to decrease with age as the oocyte quality declines (Andux and Ellis, 2008). Noteworthily, we found a reduction in the percentage of eggs that hatched upon *cyd-1* KD (**Figure 1D**). These observations indicate that *cyd-1* is required in wild-type worms to maintain the quality of germ cells and oocytes.

The *C. elegans* self-fertilizing hermaphrodite contains both sperms and oocytes. To assess if the poor oocyte quality upon *cyd-1* KD is dependent on sperm/sperm signals, we mated the *cyd-1* KD hermaphrodites to males grown on control RNAi. Interestingly, both selfed and mated hermaphrodites displayed poor oocyte morphology upon *cyd-1* depletion, implying that the poor oocyte quality is sperm-independent (**Figure 1E,F**).

Since the loss of *cyd-1* led to deterioration in oocyte quality, similar to those observed in aged worms, we asked if *cyd-1* loss accelerates the reproductive aging of WT. The reproductive span of the *cyd-1* KD worms was significantly reduced compared to control RNAi-grown worms (**Figure 1G**) with an overall reduction in the brood size (**Figure 1H**). We also observed a > 3- fold increase in the incidences of endomitotic oocytes (emos) at day-3 of adulthood in *cyd-1* KD WT worms (**Figure 1I,J**). Such uterine tumors are comparable to ovarian teratomas in humans and are often found in the uterus of aged worms, in the post-reproductive phase (Wang et al., 2018). We asked if these emos have lost their germ cell fate and acquired somatic fate, similar to teratomas. To address this, we examined the expression of *unc-119*, which is expressed pan- neuronally but not in oocytes (Maduro and Pilgrim, 1995). *cyd-1* KD resulted in *unc-119p::gfp* expressing emos (**Figure 1K,L**), thereby indicating trans-differentiation of these germ cells. We also found the presence of these densely DAPI-stained emos upon c*yd-1* KD in the feminized worms, indicating these are sperm-independent in origin (**Figure S1H**). Therefore, *cyd-1* is required to maintain the normal reproductive span and its depletion leads to accelerated reproductive aging.

### *Cyclin D/cyd-1* is required in the somatic gonad (uterus) for oocyte quality maintenance

Previously, the IIS and the TGF-β signaling in the somatic tissues (hypodermis, muscle and intestine) have been shown to regulate reproductive aging (Luo et al., 2010b). The somatic tissues crosstalk with the germline, not only to regulate aging (Berman and Kenyon, 2006; Lee et al., 2019; Yamawaki et al., 2010) but also to influence germline development (Killian and Hubbard, 2005) and the oocyte quality (Luo et al., 2010a). Keeping this in mind, we wanted to test the tissue-specific requirement of *cyd-1* in the regulation of oocyte quality. For this, we employed a tissue-specific RNAi system where in the *rde-1* (gene encoding an Argonaute protein) mutant background, a tissue-specific promoter is used to drive *rde-1* expression, providing an elegant system to knock a gene down in that tissue alone (Qadota et al., 2007). Surprisingly, in contrast to the whole-body RNAi of *cyd-1*, KD of *cyd-1* in the germ cells [using a transgenic line where *sun-1* promoter drives the expression of *rde-1* only in the germ cells of the *rde-1* mutant (Zou et al., 2019)] alone was unable to deteriorate the oocyte quality (**Figure 2A**, B), increase the percentage of endomitotic oocytes (**Figure 2C**, D) or affect the brood size (**Figure S2A**). Interestingly, somatic tissue-specific KD using a *ppw-1* [a PAZ/PIWI protein required for efficient germline RNAi (Tijsterman et al., 2002)] mutant resulted in poor oocyte quality (**Figure 2A,B**) with increased emos (**Figure 2C,D**) and a concomitant reduction in brood size (**Figure S2B**). The germline and somatic RNAi specificity of these mutants was validated by *egg-5* RNAi (Parry et al., 2009) (which leads to the laying of dead eggs when RNAi is functional in the germline) and *dpy-7* RNAi (McMahon et al., 2003) (which leads to dumpy phenotype when RNAi is functional in the soma) (**Figure S2C**).

**Figure 2.**
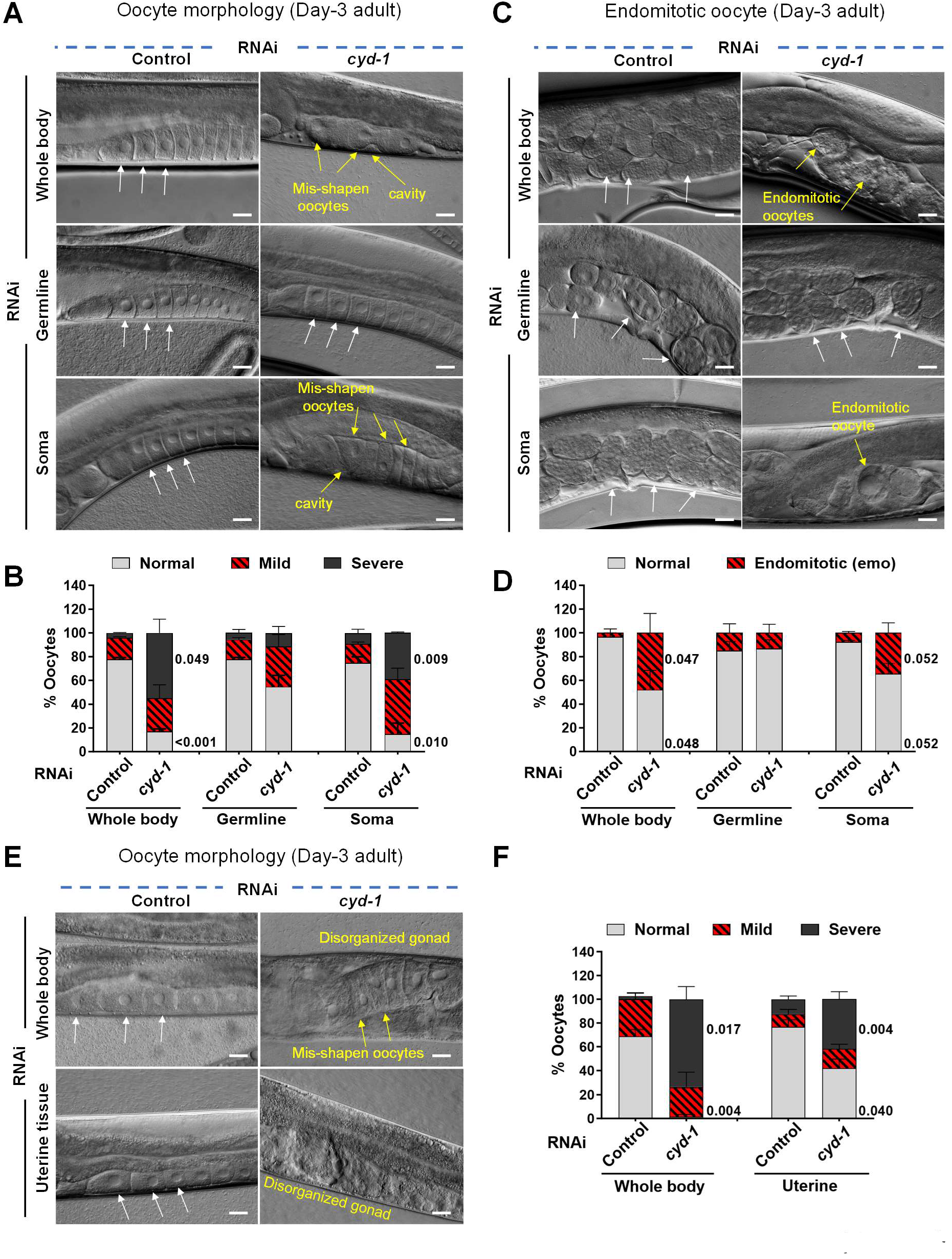
CYD-1 regulates oocyte quality cell non-autonomously from the somatic gonad (uterus) **(A,B)** DIC images showing oocyte morphology of WT, *rde-1(mkc36);sun-1p::rde-1* (germline-specific RNAi) and *ppw-1(pk1425)* (soma-specific RNAi) (day-3 adult) grown on control or *cyd-1* RNAi. White arrows- normal oocytes, yellow arrows- oocytes with abnormalities (A) Quantification of oocyte quality score (B). The quality of oocytes was categorized as normal, or mild or severe based on its morphology (cavities, shape, organization) as shown in Figure S1G. Average of three biological replicates (n ≥ 25 for each replicate). Unpaired *t*-test with Welch’s correction. **(C,D)** DIC images showing the presence of eggs or endomitotic oocytes (emos) in the uterus of WT, *rde-1(mkc36);sun-1p::rde-1* (germline-specific RNAi) and *ppw-1(pk1425)* (soma-specific RNAi) (day-3 adult) grown on control or *cyd-1* RNAi. White arrows mark normal eggs while yellow arrows mark endomitotic oocytes (emos) (C). Quantification for endomitotic oocytes (D). Average of three biological replicates (n ≥25 for each experiment). Unpaired *t*-test with Welch’s correction. **(E,F)** DIC images showing oocyte morphology of *rrf-3(pk1426)* (Whole body RNAi) or *rrf- 3(pk1426);rde-1(ne219);fos-1ap::rde-1(genomic)* (uterine tissue-specific RNAi) worms (day-3 adult) grown on control or *cyd-1* RNAi. White arrows mark normal oocytes while yellow arrows mark oocytes with severe abnormalities (E). Oocyte quality scores (based on morphology as shown in Figure S1G) (F). Average of three (whole body KD) and four (uterine specific KD) biological replicates (n ≥ 25 for each replicate). Unpaired *t*-test with Welch’s correction. Scale bars:20 μm. Error bars are s.e.m. Experiments were performed at 20°C. Source data are provided in Table S1.

To further investigate the somatic tissue where *cyd-1* may function to influence the germline quality, we used the tissue-specific strains in the *rrf-3* mutant background. Intestine or muscle-specific knock-down of *cyd-1* resulted in no changes in the oocyte morphology (**Figure S2D,E**). Surprisingly, uterine-specific KD of *cyd-1* [using a transgenic line where *fos1a* promoter drives the expression of *rde-1* only in the uterine cells of the *rde-1* mutant (Hagedorn et al., 2009)] led to compromised oocyte quality (**Figure 2E,F**) with an increase in the emos (**Figure S2F,G**) and a reduction in the total progeny number (**Figure S2H**). The strain was confirmed to have RNAi machinery specifically in the uterine cells as it showed a protruded vulva phenotype upon *egl-43* KD (Rimann and Hajnal, 2007) and no dead eggs or dumpy phenotype on *egg-5 and dpy- 7* RNAi, respectively (**Figure S2I**). Thus, *cyd-1* is important in the somatic gonad (uterine tissue) for oocyte quality maintenance in the WT germline. This exemplifies a soma-to-germline crosstalk that ensures the quality of the germline.

### *Cyclin D/cyd-1* KD leads to a DAF-16a isoform-dependent pachytene arrest in *daf-2*

While the depletion of *cyd-1* accelerates the decline in oocyte quality in WT, depletion of *cyd-1* under low insulin signaling (*daf-2* worms) leads to complete sterility, as mentioned earlier (**Figure S1A, 3A-C**). Interestingly, similar to stress resistance (Evans et al., 2008; Henderson and Johnson, 2001; McColl et al., 2010; Murphy et al., 2003) and longevity (Lin et al., 1997; Ogg et al., 1997), this germline arrest also was mediated by the downstream FOXO/DAF-16 TF, such that the fertility is rescued in *the daf-16;daf-2;cyd-1* RNAi worms (**Figure 3A-C**). To determine the effect of *cyd-1* KD on germline development of *daf-2*, the gonads of day-1 adults were stained with DAPI. Interestingly, the *cyd-1* KD did not lead to a reduction in mitotic germ cell number. Importantly, however, no oocytes were formed upon *cyd-1* KD in *daf-2* and the oocyte formation was restored in the *daf-16;daf-2* double mutants (**Figure 3D,E**). Upon quantification of the germ cell numbers, we found that *cyd-1* KD led to small but significant changes in the transition zone germ cells, but drastically reduced the pachytene germ cell number in *daf-2*; this was restored in *daf-16;daf-2* (**Figure 3D,E**). We measured *cyd-1* mRNA levels and confirmed similar efficiency of *cyd-1* KD in *daf-2* and *daf-16;daf-2* worms, implying that the germline arrest is not due to a differential *cyd-1* KD efficiency (**Figure S3A**). We further speculated if KD of *cyd-1* in *daf-2* worms resulted in critically low levels of *cyd-1* levels below a presumptive threshold, thereby causing sterility. To rule out this, we KD *cyd-1* by RNAi in a balanced *cyd-1* mutant, *cyd-1(he112)* that contains a truncated C-terminal due to the presence of a nonsense mutation (Boxem and van den Heuvel, 2001). Interestingly, we observed deterioration of oocyte quality but no sterility, implying the arrest is specific to *daf-2* (data not shown). Thus, upon depletion of *cyd-1* in *daf-2*, activated DAF-16 plays a decisive role in arresting the germ cells at the pachytene stage of meiosis and in preventing oocyte formation. One of the causes of sterility in *daf-2;cyd-1* RNAi worms could be elevated levels of germ cell apoptosis. For measuring apoptosis, we employed the *ced-1::gfp* transgenic strain as described above. Elevated levels of apoptosis were observed for both *daf-2* and *daf-16;daf-2* worms grown on *cyd-1* RNAi (**Figure S3B,C**), implying that the sterility in *daf-2* upon *cyd-1* KD may not be solely due to an increase in apoptosis. We wondered if the oocytes that bypass the pachytene arrest in *daf-16;daf-2;cyd-1* RNAi worms would also show hallmarks of reproductive aging. Indeed, we observed that *cyd-1* KD in *daf-16;daf-2* led to severe defects in oocyte quality (**Figure S3D,E**) as well as an increased occurrence of emo in the uterus of these animals (**Figure S3F,G**).

**Figure 3.**
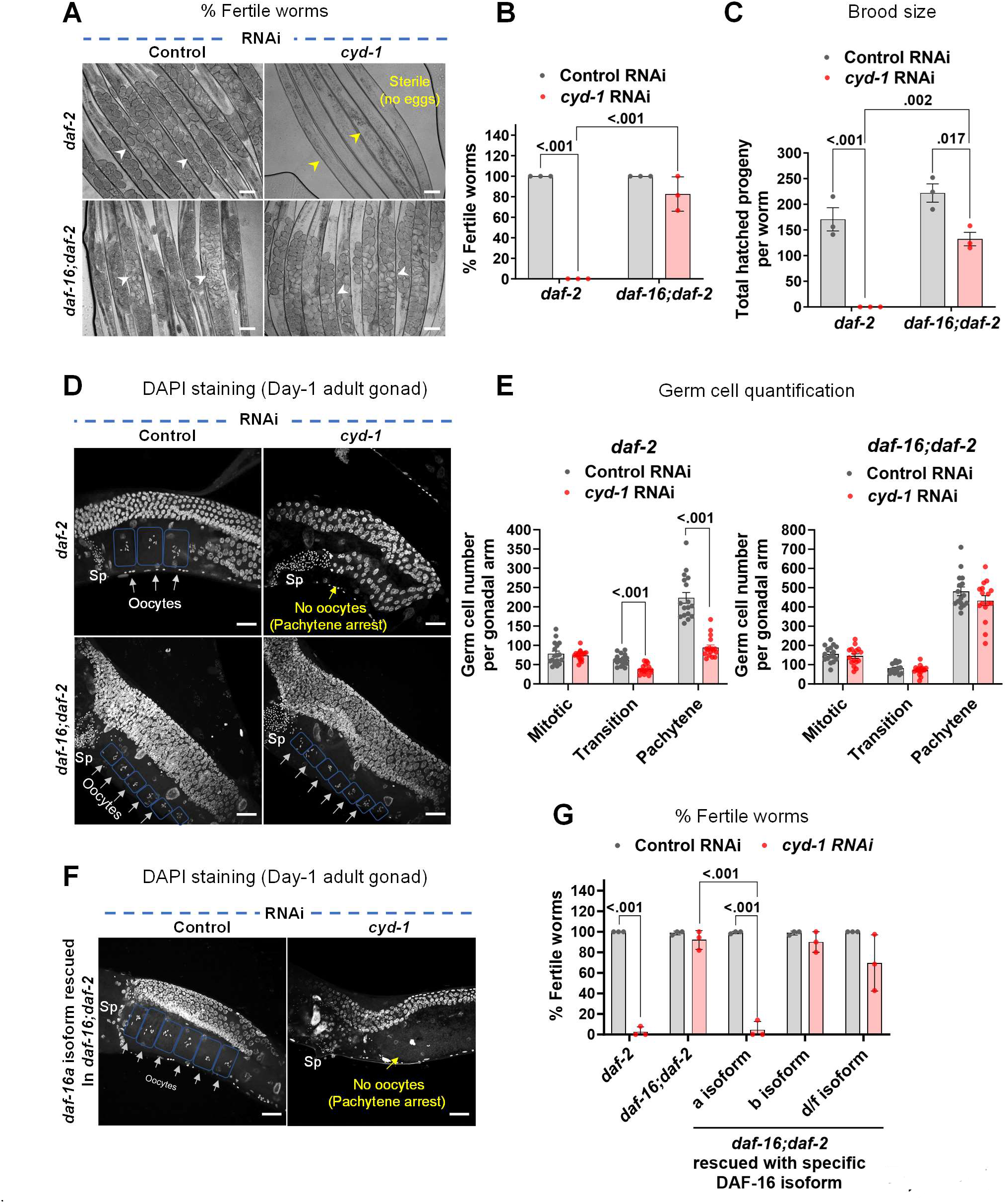
*cyd-1* KD leads to FOXO/DAF-16-dependent germline arrest under low insulin signaling. **(A,B)** Representative images showing that *cyd-1* RNAi results in sterility in *daf-2(e1370)* worms that are rescued in *daf-16(mgdf50);daf-2(e1370)*. White arrowheads show eggs while yellow arrowheads show gonads with no eggs (sterile worms) (A). The percentage of fertile worms (B). Average of three biological replicates (n ≥ 30 for each experiment). Two-way ANOVA-Tukey’s multiple comparisons test. **(C)** The total number of hatched progenies in *daf-2(e1370)* and *daf-16(mgdf50);daf-2(e1370)* worms grown on control or *cyd-1* RNAi. Average of three biological repeats. Two-way ANOVA-Tukey’s multiple comparisons test. **(D,E)** Representative fluorescent images of DAPI-stained gonads of *daf-2(e1370)* and *daf- 16(mgdf50);daf-2(e1370)* (day-1 adult) worms grown on control or *cyd-1* RNAi. Oocytes are boxed for clarity. White arrows point towards oocytes while yellow shows the absence of oocytes. Sp denotes sperms (D). Quantification of DAPI-stained germ cell nuclei. n=17 (*daf-2*); n=16 (*daf- 16;daf-2*). (E) Each point represents the number of mitotic (MT), transition (TS) or pachytene zone (PZ) cells. Unpaired *t*-test with Welch’s correction. **(F,G)** Isoform requirement of DAF-16 to mediate germline arrest in *daf-2(e1370)*. Representative fluorescent images of DAPI-stained germ line of *daf-16(mgdf50);daf-2(e1370);daf-16a(+)* worms grown on control or *cyd-1* RNAi. Oocytes are boxed for clarity. White arrows point towards oocytes while yellow shows the absence of oocytes. Sp denotes sperms (F). Percentage of the fertile worms (G). The *cyd-1* was knocked down in *daf-2(e1370), daf-16(mgdf50);daf-2(e1370)* as well as in strains where different DAF-16 isoforms were rescued in *daf-16(mgdf50);daf-2(e1370).* Average of three biological replicates (n ≥ 30 for each replicate). Two-way ANOVA-Tukey’s multiple comparisons test. Scale bars:20 μm. Error bars are s.e.m. Experiments were performed at 20°C. Source data are provided in Table S1.

Different isoforms of DAF-16 exist due to alternative splicing events as well as alternate promoter usage. The different isoforms have distinct tissue-specific expression profiles and the long life span of *daf-2* depends on each isoform to a different extent (Kwon et al., 2010). We next asked which isoform of DAF-16 was necessary for inducing arrest in *daf-2* following KD of *cyd-1*. To address this, we used transgenic strains, wherein, different isoforms of DAF-16 were rescued in the *daf-16;daf-2* double mutants (Kwon et al., 2010). Interestingly, we found that the *cyd-1* KD in the *daf-16;daf-2;daf-16a(+)* strain led to sterility comparable to the *daf-2* (**Figure 3F,G**), signifying a predominant role of DAF-16a isoform in mediating this germline pachytene arrest.

Additionally, the sterility due to *cyd-1* KD in *daf-2* involves the canonical IIS pathway components as a similar arrest at the pachytene stage of meiosis was observed upon *cyd-1* KD in *age-1(hx546)* [mutant in mammalian PI3K ortholog (Dorman et al., 1995; Friedman and Johnson, 1988)] and *pdk-1(sa680)* [mutant in mammalian PDK ortholog (Paradis et al., 1999)] (**Figure S3H,I**). Interestingly, KD of the canonical binding partner of *cyd-1*, *cdk-4* (Kato et al., 1993) did not lead to sterility in *daf-2* even in the RNAi hypersensitive background (**Figure S3J**), hinting towards a cyclin-independent function of *cyd-1* in oogenesis.

### *Cyd-1* KD in the somatic gonad of *daf-2* results in germline arrest at the pachytene stage

Since we found a role for somatic *cyd-1* in regulating WT germline aging, we asked if somatic *cyd-1* KD would lead to germline arrest in *daf-2*. We first checked if the germline arrest was mediated cell-autonomously by *cyd-1* depletion in the germ cells. Interestingly, *cyd-1* depletion in the germ cells alone did not lead to sterility in *daf-2* (**Figure 4A**, B). However, *cyd-1* depletion specifically in the uterine cells *per se* resulted in the germ cell arrest at the pachytene stage (**Figure 4A**, B) and also caused a reduction in the pachytene germ cell number (**Figure 4C**). This somatic requirement of *cyd-1* in germline arrest reiterates a non-canonical role of *cyd- 1* in oogenesis. Loss of *cyd-1* in other somatic tissues like muscle, neurons, intestine, hypodermis, Distal Tip Cell (DTC) and the vulva did not cause sterility in *daf-2* (**Figure S4A**). Interestingly, we observed vulval defects (protruded vulva/vulvaless) upon *cyd-1* depletion in *daf-2* while *daf- 16;daf-2* showed no such defects (**Figure S4B,C**).

**Figure 4.**
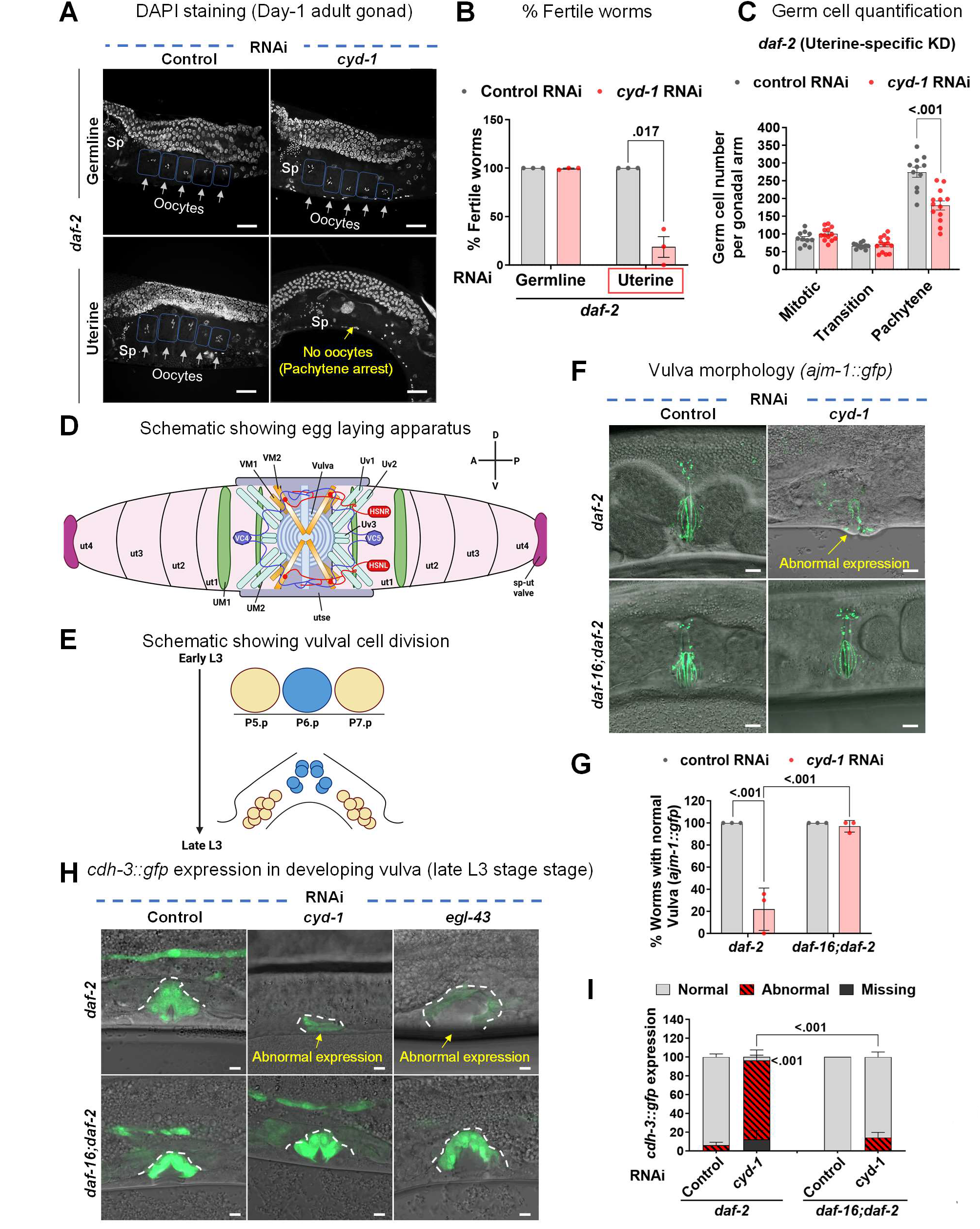
*cyd-1* KD in the somatic gonad (uterus) alone is sufficient to cause germline arrest under low insulin signaling. **(A,B)** Representative images of DAPI-stained gonads showing no pachytene arrest in day-1 adult of *daf-2(e1370);rde-1(mkc36);sun-1p::rde-1* (germline-specific RNAi) worms upon *cyd-1* KD. However, pachytene arrest ensues upon *cyd-1* KD in *daf-2(e1370);rde-1(ne219);fos-1ap::rde-1* (uterine tissue-specific RNAi). Oocytes are boxed for clarity. White arrows point towards oocytes while yellow shows the absence of oocytes. Sp denotes sperms. Scale bar 20 µm (A). The percentage of fertile worms upon germline-specific and uterine-specific *cyd-1* KD in *daf-2(e1370)* (B). Average of three biological replicates (n ≥ 25 per condition for each experiment). Unpaired *t*- test with Welch’s correction. **(C)** Quantification of DAPI-stained *daf-2(e1370);rde-1(ne219);fos-1ap::rde-1* (uterine tissue-specific RNAi) (day-1 adult) germ cell nuclei of different stages (mitotic (MT), transition (TS) or pachytene zones (PZ)); n=11. Each point represents the number of mitotic (MT), transition (TS) or pachytene zone (PZ) cells. Unpaired *t*-test with Welch’s correction. **(D)** Schematic showing the egg laying apparatus of an adult *C. elegans* hermaphrodite. The egg-laying apparatus consists of the uterus, the uterine muscles, the vulva, the vulval muscles, and the egg-laying neurons. A stack of seven nonequivalent epithelial toroids (rings) form the vulva. The anterior and posterior lobes of uterus each comprises of four uterine toroid epithelial cells (ut1–ut4). Adherens junctions connect the neighboring uterine toroids or vulval toroids, respectively. The uterine seam cell (utse) is a H shaped cell that connects the uterus to seam cells and thus holds the uterus in place. uv1-3 are interfacial cells between uterus and vulva (uv1 is a neuroendocrine cell). The process of egg-laying is made possible through the contraction of sex muscles, namely the vulval muscles (VM1,2) connected to the vulva lips and the uterine muscles (UM1,2) encircling the uterus. The activity of these muscles is under the regulation of motor neurons, specifically VCn (VC1-6) and HSNL/R, which form synapses with each other and with the vulval muscle arms. A=anterior, P=posterior, D=dorsal, V=ventral. Adapted from wormbook (doi:10.3908/wormatlas.1.24). **(E)** Schematic showing the vulval cell divisions. The vulval precursor cells (VPCs) P5.p, P6.p and P7.p are induced to divide in the early L3 stage. Three rounds of cell division yield a total of 22 vulval cells by the end of L3 stage. The adult vulva is then formed through subsequent patterning and morphogenesis processes. **(F,G)** Representative fluorescent and DIC merged images of gonads showing *ajm-1::gfp* (marks cell junctions) expression in *daf-2(e1370)* and *daf-16(mgdf50);daf-2(e1370)* worms (day-1 adult) grown on control and *cyd-1* RNAi. Yellow arrows point towards abnormal expression. Scale bar 10 µm (F). Quantification of the normal or abnormal expression of *ajm-1::gfp* (G). Average of three biological replicates (n ≥ 24 for each experiment). Two-way ANOVA-Tukey’s multiple comparisons test. **(H,I)** Representative fluorescent and DIC merged images of gonads showing *cdh-3::gfp* expression in *daf-2(e1370)* and *daf-16(mgdf50);daf-2(e1370)* worms (late-L3 stage) on control, *cyd-1* or *egl-43* RNAi. Yellow arrows point towards abnormal expression. Scale bar 5 µm (H). Quantification for normal, reduced or missing *cdh-3::gfp* expression (I). Average of three biological replicates (n ≥ 30 for each experiment). Two-way ANOVA-Tukey’s multiple comparisons test. Error bars are s.e.m. Experiments were performed at 20°C. Source data are provided in Table S1.

The egg-laying apparatus consists of the uterus, the uterine muscles, the vulva, the vulval muscles, and the egg-laying neurons. The uterus and vulva are connected by the uterine seam cell (Newman et al., 2000) (**Figure 4D**). The vulva formation starts at the L3 larval stage and vulval cell-divisions are completed by the late-L3 stage (**Figure 4E**). We investigated if the loss of *cyd-1* caused defects in the vulva-uterine morphology. We visualized the vulval muscle architecture by Phalloidin staining (that marks the F-actin) that revealed a distorted vulva muscle structure following the removal of *cyd-1* in *daf-2* (**Figure S4D**). To further dissect the underlying cause of the vulval defects, we employed the *ajm-1::gfp* transgenic strain (which marks the adherens junction) (Koppen et al., 2001). Vulval development involves the formation of a tubular structure from only 22 epithelial cells. It encompasses an array of steps including vulval cell induction and differentiation, cell division, cell rearrangements, invagination, toroid formation and cell-cell fusion and attachment with other tissues (hypodermis, muscle, uterus) (Gupta et al., 2012). Normally, in the vulva, the AJM-1::GFP marks the symmetrical toroid; however, upon *cyd- 1* depletion in *daf-2*, the AJM-1::GFP was mis-localized and the expression was diminished. In contrast, loss of *cyd-1* in *daf-16;daf-2* worms did not exhibit such abnormalities (**Figure 4F,G**). This highlights the crucial role of activated DAF-16 in causing vulval impairments consequent to *cyd-1* KD. Next, we asked if these defects in vulva were due to impairment in the vulval cell divisions. We observed the vulval precursor cell (VPC) fate of late-L3 staged worms using the transgenic reporter strain, *cdh-3::gfp* (Pettitt et al., 1996). In *daf-2* worms fed control RNAi, we observed the correct expression of *cdh-3::gfp* in vulval cells, however, the expression was abnormal or absent upon *cyd-1* RNAi, suggesting incomplete or aberrant vulval cell divisions.

Notably, the *daf-16;daf-2;cyd-1* RNAi animals exhibited a restoration of normal *cdh-3::gfp* expression in vulval cells (**Figure 4H,I**). Furthermore, we utilized the depletion of the *egl-43* gene, essential for vulval morphogenesis (Hwang et al., 2007; Rimann and Hajnal, 2007), as a control, and observed analogous abnormalities in *cdh-3::gfp* expression. Interestingly, in this context as well, we observed a DAF-16-dependence such that the *cdh-3::gfp* expression was restored in the *daf-16;daf-2* worms (**Figure 4H**). Additionally, knockdown of *egl-43* in *daf-2* resulted in endomitotic oocytes and sterility, which was partially rescued in the *daf-16;daf-2* worms (**Figure S4E,F)**. These observations allude to the possibility that activated DAF-16 may amplify deficiencies in the egg-laying apparatus.

### *Cyd-1* KD results in DAF-16-dependent defects in the somatic gonad of *daf-2* worms

To gain deeper insights into the changes underlying the germline arrest due to *cyd-1* KD in *daf-2*, we performed transcriptomics analysis of the late-L4 staged WT and *daf-2* worms with *cyd-1* KD only in the gonad (using a transgenic strain where *Punc-52* drives the expression of *rde-1* in the *rde-1* mutant). Interestingly, the genes important for cell cycle/cell-fate, oogenesis and reproduction were significantly downregulated in *daf-2;cyd-1* RNAi worms (**Figure S5A**), but not in *cyd-1* KD in the WT worms (**Figure S5B**). Importantly, the genes involved in vulva-uterine [*lin-11* (Newman et al., 1999)*, lim-6* (Hobert et al., 1999), *unc-59* (Nguyen et al., 2000)] and sheath cell development [*lim-7*(Voutev et al., 2009)] were also down-regulated in the *daf-2* gonad-specific *cyd-1* KD (**Figure 5A**, Table S2) but not in WT (**Figure S5C**, Table S3). The gonadal sheath cells are a crucial component of the worm somatic gonad that plays an important role in pachytene exit and oogenesis (Govindan et al., 2006; Hall et al., 1999; McCarter et al., 1997b; Whitten and Miller, 2007). The 5 pairs of sheath cells surround each gonadal arm (**Figure 5B**). We tested if *cyd-1* KD could also lead to defects in the sheath cells. For this, we used the sheath cell marker strain (*lim- 7p::gfp*) (Hall et al., 1999) in the WT, *daf-2* or *daf-16;daf-2* worms. Interestingly, we found that the *cyd-1* KD in *daf-2* led to the complete abrogation of the *lim-7p::gfp* expression, which was restored in the *daf-16;daf-2* (**Figure 5C,D**). We stained the gonads of day-1 adults with phalloidin (labels F-actin) to investigate potential alterations in sheath cell structure in *daf-2;cyd-1* RNAi worms. Strikingly, we observed a significant disruption in the cytoskeletal organization of sheath cells in *daf-2;cyd-1* RNAi worms, which was ameliorated in *daf-16;daf-2* worms subjected to *cyd-1* RNAi (**Figure 5E,F,S5D**). We inquired if KD of genes essential for sheath cell development (eg. *lim-7, lim-6)* would also lead to germline arrest in *daf-2* but found no such sterility (data not shown). However, similar to *cyd-1* RNAi we observed DAF-16-dependent pachytene arrest and sterility (**Figure 5G,H**) and repression of *lim-7p::gfp* expression (**Figure S5E,F**) upon KD of the *sys-1/β-catenin* transcription factor downstream of WNT signaling, which is known to play a key role in the development of the somatic gonad (Phillips et al., 2007; Siegfried et al., 2004). This suggests that a more drastic somatic gonad developmental defect is required to cause DAF-16-dependent sterility in *daf-2*.

**Figure 5.**
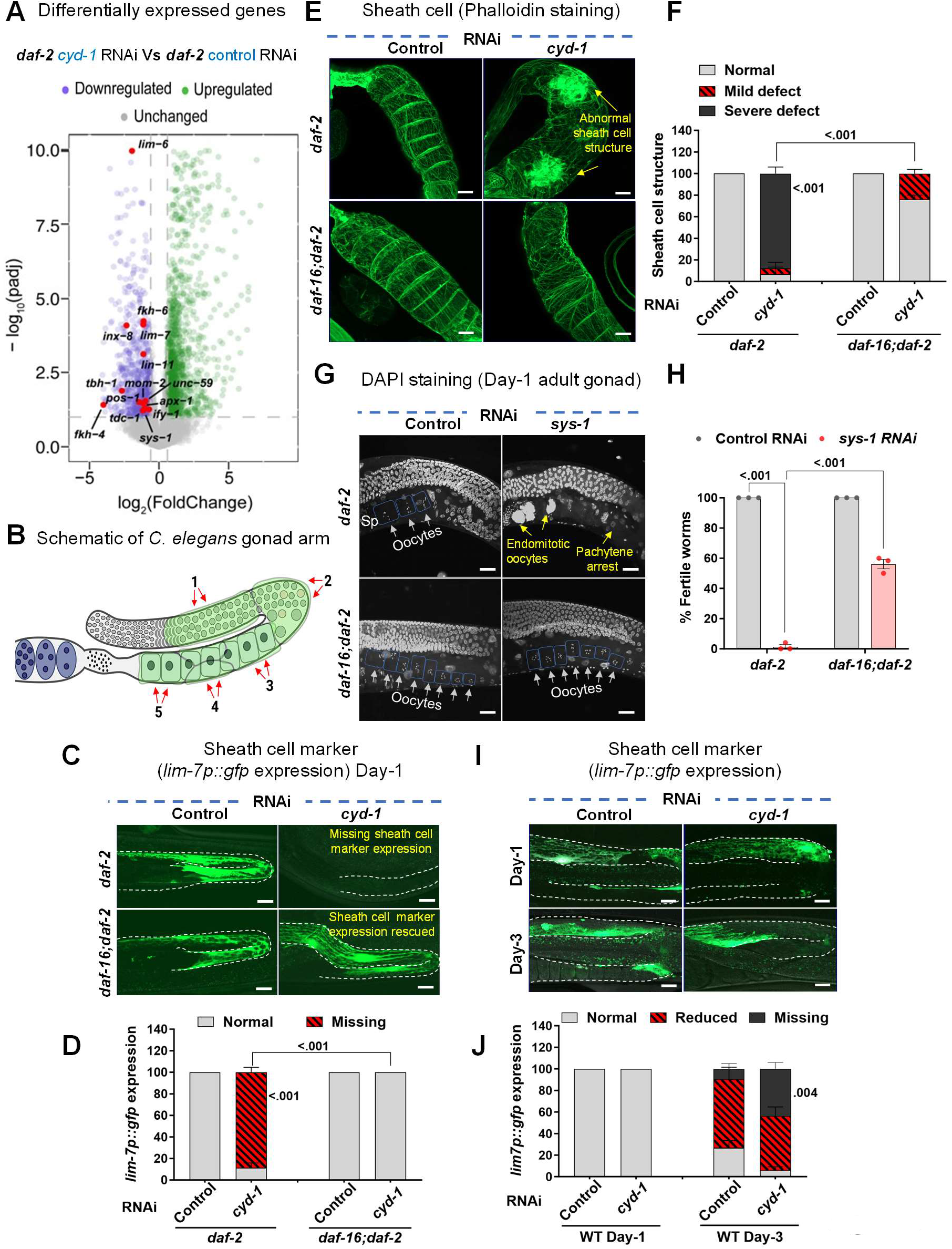
Loss of *cyd-1* under low insulin signaling leads to DAF-16 dependent defects in sheath cells. **(A)** Volcano plot showing the magnitude [log2(FC)] and significance [−log10(P value)] of the genes that are differentially expressed in L4 staged *daf-2(e1370);rrf-3(pk1426);rde-1(ne219);unc- 62p::rde-1(genomic)* worms, grown on control or *cyd-1* RNAi. Y axis [−log10(P value)] is restricted to a value of 10. **(B)** Schematic showing one of the two gonad arms of an adult *C. elegans* hermaphrodite. Sheath cells are colored in green. **(C,D)** Representative fluorescent and DIC merged images of gonads showing *lim-7p::gfp* (that marks the sheath cells) expression in *daf-2(e1370)* and *daf-16(mgdf50);daf-2(e1370)* worms (day-1 adult) grown on control and *cyd-1* RNAi. The gonadal arm is outlined for clarity. Scale bar 20 µm (C). Quantification of the normal, reduced or missing expression of *lim-7p::gfp* (D). Average of three biological replicates (n ≥ 18 for each experiment). Two-way ANOVA-Tukey’s multiple comparisons test. **(E,F)** Representative fluorescent images of phalloidin-stained (that marks the F-actin) gonads of *daf-2(e1370)* and *daf-16(mgdf50);daf-2(e1370)* worms (day-1 adult) grown on control and *cyd-1* RNAi. Arrows point towards defects in the sheath cells. Scale bar 10 µm (E). Quantification of the normal or defective sheath cell structure (as per scoring scheme in Figure S5D) (F). Average of three biological replicates (n ≥ 13 for each experiment). Two-way ANOVA-Tukey’s multiple comparisons test. **(G,H)** Representative fluorescent images of DAPI-stained gonads of *daf-2(e1370)* and *daf- 16(mgdf50);daf-2(e1370)* (day-1 adult) worms grown on control or *sys-1* RNAi. Oocytes are boxed for clarity. White arrows point towards oocytes while yellow shows endomitotic oocytes or lack of oocytes. Scale bar 20 µm (G). The percentage of fertile worms (H). Average of three biological replicates (n ≥ 25 per condition for each experiment). Two-way ANOVA-Tukey’s multiple comparisons test. **(I,J)** Representative fluorescent images of gonads showing *lim-7p::gfp* (that marks the sheath cells) expression in WT worms (day-1 and day-3 adult) grown on control and *cyd-1* RNAi. Arrows point towards defective/missing *lim-7p::gfp* expression in the sheath cells. The gonadal arm is outlined for clarity. Scale bar 20 µm (I). Quantification of the normal/defective or missing *lim-7:: gfp* expression (J). Average of three biological replicates (n ≥ 18 for each experiment). Unpaired *t*-test with Welch’s correction. Error bars are s.e.m. Experiments were performed at 20°C. Source data are provided in Table S1.

Next, we investigated if *cyd-1* RNAi can alter the expression of another adult hermaphrodite gonad-specific marker (*fkh-6::gfp*). In the hermaphrodite gonad, FKH-6 is expressed in the adult spermatheca (Chang et al., 2004). Similar to *lim-7p::gfp* expression, we found a DAF-16-dependent suppression of *fkh-6::gfp* expression upon *cyd-1* depletion in *daf-2* (**Figure S5G,H**). Interestingly, while the WT worms showed no obvious structural defects in the sheath cell upon *cyd-1* depletion (**Figure S5I,J**), *lim-7p::gfp* expression was significantly reduced in day-3 *cyd-1* KD WT worms compared to age-matched control worms (no difference in day-1 adults) (**Figure 5I,J**).

Overall, these data imply that loss of *cyd-1* elicits a DAF-16-dependent distortion of sheath cell structure and suppression of hermaphrodite gonad marker expression in *daf-2*. The amplification of somatic gonadal abnormalities by activated DAF-16 may, in turn, influence oogenesis.

### *Cyd-1* KD results in a defective spermatogenesis-to-oogenesis fate switch in *daf-2*

In the hermaphrodites, a fixed number of sperms are formed in the L4 larval stage, after which the fate is switched from spermatogenesis-to-oogenesis and adult worms make only oocytes from then onwards (Ahringer and Kimble, 1991). Upon knockdown of *cyd-1* in *daf-2*, we observed only sperms but no oocytes in the adult hermaphrodite worms (**Figure 6A**). Moreover, we showed above that lowering *cyd-1* levels in *daf-2* suppresses the hermaphrodite gonad-specific marker expression (**Figure 5C,D,S5G,H**). We supposed that either the germline has been masculinized (male mode of germline: continuous sperm production) or there is a failure in the spermatogenesis-to-oogenesis fate switch. However, we did not observe any ectopic expression of the male gonad marker (*K09C8.2p::gfp)* (Thoemke et al., 2005) in *daf-2;cyd-1* RNAi worms (**Figure S6A**). Also, the sperm number remained unchanged in the *daf-2* young adult with *cyd-1* KD (**Figure 6B**) and the sperm number did not increase on day-2 or day-3 of adulthood, indicating that the germline had not been masculinized (**Figure 6C**). Alternatively, *cyd-1* loss in *daf-2* may cause a defect in the spermatogenic-to-oogenic cell fate switch and a consequent arrest of oogenesis at the pachytene stage. In line with this assumption, the depletion of *cyd-1* led to a DAF-16-dependent downregulation of genes essential for the sperm-to-oocyte fate switch (**Figure 6D,E**). Since the proximal-most sheath cell pairs play an important role in the pachytene exit (Killian and Hubbard, 2005; McCarter et al., 1997a), the deformation of sheath cells in the *daf- 2;cyd-1* RNAi worms may be the driver for the defective sperm-to-oocyte fate switch, leading to sterility in these animals.

**Figure 6.**
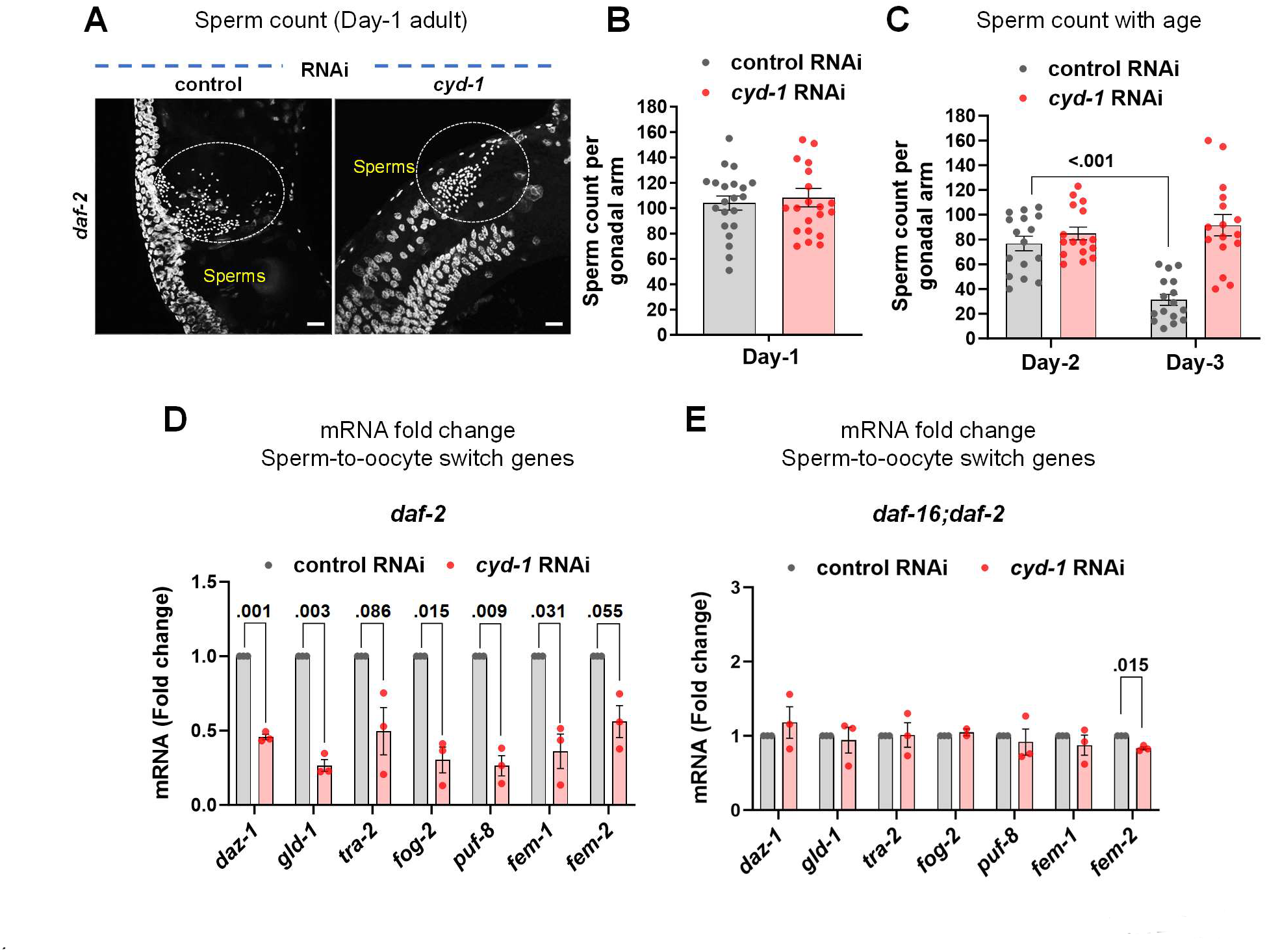
A defective spermatogenesis-to-oogenesis switch underlies germline arrest upon *cyd-1* KD under low insulin signaling. **(A,B)** Representative fluorescent images of DAPI stained *daf-2(e1370)* worms (early day-1 adult) grown on control and *cyd-1* RNAi. Sperms are encircled for clarity. Scale bar 10 µm (A). Quantification of the sperm count (B).n = 21 for each condition. Each point represents the number of sperms per gonadal arm per worm. Unpaired *t*-test with Welch’s correction. **(C)** Quantification of the sperm count in *daf-2(e1370)* worms (day-2 and day-3 adults) grown on control and *cyd-1* RNAi. n =16 for each condition. Each point represents the number of sperms per gonadal arm per worm. Two-way ANOVA-Tukey’s multiple comparisons test. **(D)** Quantitative RT-PCR analysis of sperm-to-oocyte switch genes in *daf-2(e1370)* worms (late- L4 stage) grown on control or *cyd-1* RNAi. Expression levels were normalized to *actin*. Average of three biological replicates are shown. Unpaired *t*-test with Welch’s correction. **(E)** Quantitative RT-PCR analysis of sperm-to-oocyte switch genes in *daf-16(mgdf50);daf- 2(e1370)* worms (late-L4 stage) grown on control or *cyd-1* RNAi. Expression levels were normalized to *actin*. Average of three biological replicates are shown. Unpaired *t*-test with Welch’s correction. Error bars are s.e.m. Experiments were performed at 20°C. Source data are provided in Table S1.

## Discussion

The role of Cyclin D/CDK-4 in the progression of G1 to S phase of the cell cycle is well known. Research in the past decade has shed light on several non-cell cycle functions of cyclin D, including its role in adipogenesis (Abella et al., 2005; Phelps and Xiong, 1998), muscle differentiation (Cenciarelli et al., 1999; Kiess et al., 1995), anoikis (Gan et al., 2009), DNA double-strand break (DSB) repair (Jirawatnotai et al., 2011) and transcriptional regulation (Adnane et al., 1999; Iwatani et al., 2010; Ratineau et al., 2002), many of them independent of its binding partner, CDK-4 (Inoue and Sherr, 1998; Jian et al., 2005; Jirawatnotai et al., 2011; Yu et al., 2013). Here, we decipher a novel role of *C. elegans cyclin D/cyd-1* in preserving the quality of oocytes. Depletion of *cyd-1* in wild-type nematodes led to misshapen oocytes with increased cavities between them and disruption of gonadal organization, akin to that of aged worms. Concurrently, endomitotic oocytes increased in the germline of these worms, with a decline in egg hatchability and reproductive span (**Figure 7**). These findings underscore the significance of *cyclin D* in the regulation of reproductive aging. At an organismal level, environmental cues are integrated by the nutrient sensing pathways that often coordinate with cell-cycle regulators to influence reproductive cell fate (Eustice et al., 2022; Fukuyama et al., 2012; Korta et al., 2012; Narbonne and Roy, 2006). Lowering a key nutrient-sensing pathway, the worm IIS, causes activation of the downstream FOXO/DAF-16 transcription factor leading to improved oocyte quality and delayed reproductive senescence (Hughes et al., 2007; Luo et al., 2010a). In mammals too, FOXO3a has been shown to maintain the ovarian reserve (Pelosi et al., 2013; Tejerina Botella et al., 1992), highlighting the conserved role of FOXO in safeguarding oocyte health. In this study, loss of *cyd-1* in the *daf-2* worms (low IIS) resulted in a FOXO/DAF-16-dependent sterility due to the arrest of germ cells at the pachytene stage of meiosis-I. The canonical components of the IIS/PI3K pathway, with a predominant role of DAF-16a isoform, were found to be required to mediate this germline arrest. Noteworthily, the oocytes of *daf-16;daf-2* mutants that escape the pachytene arrest induced by *cyd-1* KD exhibit poor quality, highlighting the importance of DAF-16 in sensing perturbations in the cell cycle and halting oogenesis, thereby preventing the production of unhealthy progenies (**Figure 7**). The role of FOXO proteins has been well documented as a checkpoint, where in various cell line models it orchestrates cell-cycle arrest and repair in response to DNA damage and oxidative stress (Furukawa-Hibi et al., 2002; Hill et al., 2014; Hornsveld et al., 2021; Tran et al., 2002). We have earlier shown that DAF-16 responds to somatic DNA damage and arrests germline to protect the progeny (Sarkar et al., 2023). Also, on sensing the unavailability of food, newly hatched worms utilize DAF-16 to arrest at the L1 diapause stage and resume development upon re-feeding (Baugh and Sternberg, 2006; Jensen et al., 2006). Similarly, DAF-16 has a crucial role in another well-studied alternative third larval diapause stage; dauer, that forms in response to high population density or low food availability (Gottlieb and Ruvkun, 1994). In both the above instances, crowding and low food availability would potentially jeopardize the success of the next generation. Thus, these instances reinforce the role of FOXO/DAF-16 as a custodian of both somatic as well as reproductive fitness, ultimately facilitating the efficient perpetuation of the genome.

**Figure 7.**
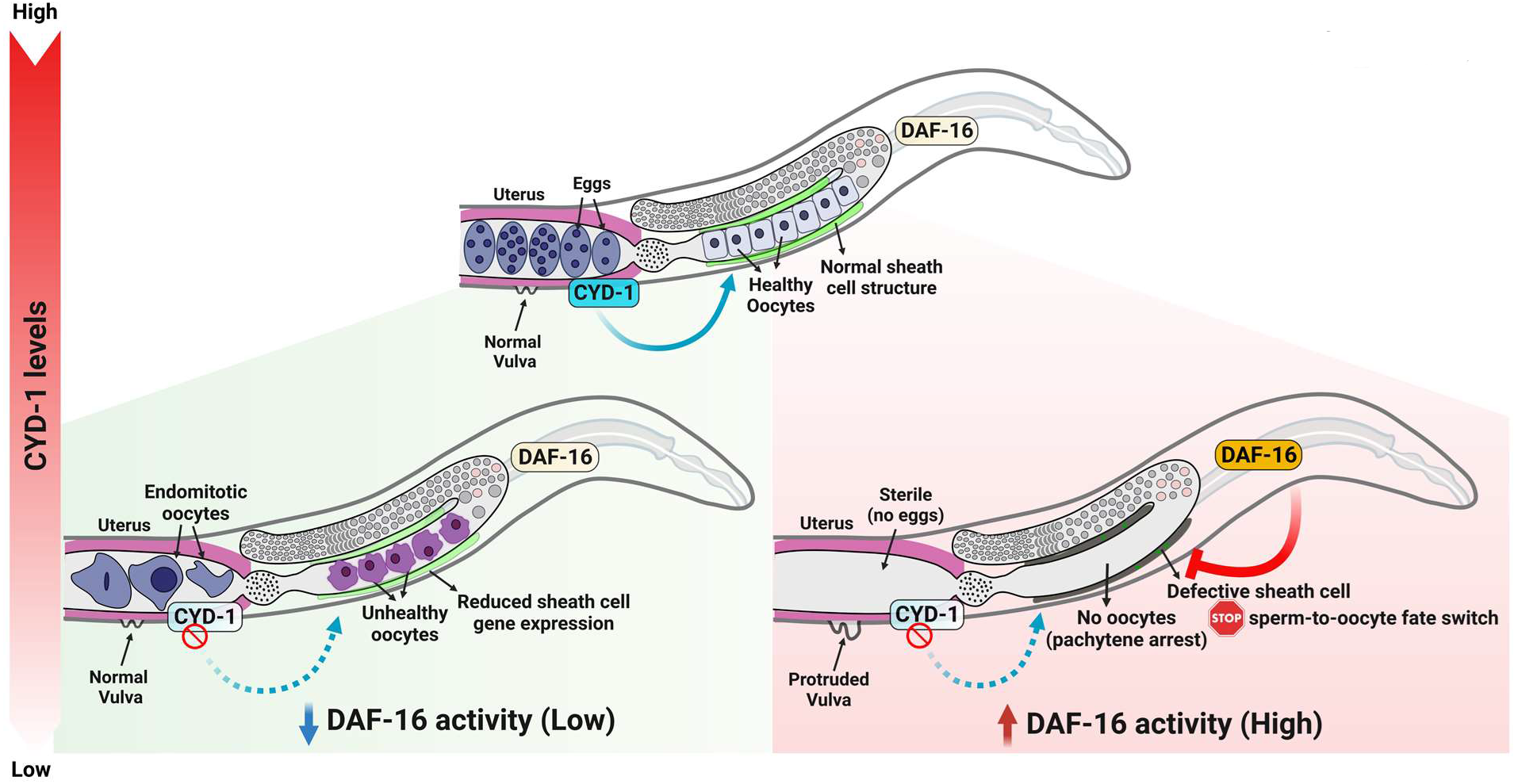
Summary model. **(A)** Model showing combined effect of *cyd-1* levels and DAF-16 activation state on the oogenesis and oocyte quality. Under wild-type conditions (normal insulin signaling) CYD-1 is required to maintain the oocyte health. Loss of *cyd-1*, leads to compromise in oocyte quality and an increase in endomitotic oocytes in the uterus. Interestingly, when DAF-16 is activated (in *daf-2* worms), loss of *cyd-1* results in DAF-16-dependent sheath cell defects leading to failure in the sperm-to-oocyte fate switch and halting of oogenesis (pachytene arrest).

Inter-tissue communication ensures an effective integration of environmental cues for optimal reproductive decisions. For example, activation of DAF-16 is required in the muscle and intestine for oocyte quality maintenance (Luo et al., 2010a) and in the uterus for maintenance of germline stem cell (GSC) pool with age (Qin and Hubbard, 2015). In line with this, we also find that *cyd-1* regulates oocyte quality and pachytene arrest of the germ cells, cell non-autonomously from the somatic gonad (uterus) under wild-type and low IIS, respectively (**Figure 7**). We have previously demonstrated that somatic DNA damage triggers DAF-16/FOXO-dependent stalling of oogenesis under low IIS in a cell non-autonomous manner (Sarkar et al., 2023). In a different context, somatic ERK/MPK-1 (Robinson-Thiewes et al., 2021) and *kri-1* (Ito et al., 2010) have been shown to influence germ cell proliferation and death, respectively. The soma-germ cell communication is also vital for mammalian reproductive competence where the oocyte and surrounding somatic granulosa cells (GC) and the cumulus cells (CC) crosstalk to coordinate the growth and development of the follicle (Norris et al., 2009; Park et al., 2004; Robinson et al., 2012; Zuccotti et al., 2011).

Intriguingly, contrary to the widely appreciated pro-longevity role of DAF-16, we observed activated DAF-16-dependent disruption of the somatic gonad morphology and gene expression upon *cyd-1* depletion under low IIS. We noticed abnormalities in the vulva (protruding vulva/vulvaless) and defects in the surrounding gonadal sheath cell structure. Consistent with this, we found transcriptional down-regulation of many genes important for somatic gonad development upon loss of *cyd-1* in *daf-2* worms, in a DAF-16-dependent manner (**Figure 7**). Activated DAF-16 led to complete abrogation of the hermaphrodite sheath cell-specific *lim-7p::gfp* and spermatheca-specific *fkh-6::gfp* expression in the *cyd-1*-depleted *daf-2* worms; both the structural defects in the gonadal sheath cells as well as gene expression alterations were reversed in the *cyd-1-*depleted *daf-16;daf-2* worms. Consistent with the role of sheath cells in the sperm-to-oocyte fate switch, we observed the downregulation of genes crucial for the sperm-to- oocyte switch in *cyd-1-*depleted *daf-2* worms. Notably, the removal of DAF-16 in these worms restored the transcript levels of these genes to normal levels, enabling the pachytene exit of oogenic germ cells (**Figure 7**). At first, this appears strikingly counter-intuitive to the traditionally recognized role of FOXO/DAF-16 in the maintenance of somatic and reproductive health. However, FOXO/DAF-16 is well-established as a sentinel, often arresting the cells or halting development on encountering stresses such as starvation, DNA damage or oxidative stress, only to provide time for damage repair or resume growth/development upon return of favorable environmental conditions. Therefore, we speculate that activated DAF-16 may sense the perturbations in the cell cycle (on *cyd-1* KD) and exacerbate defects in the somatic gonad, driving a failure in the sperm-to-oocyte fate switch as a means of halting the production of poor-quality oocytes, thereby ensuring progeny fitness. Defects in cell cycle proteins in the somatic tissue may be interpreted as a potential threat to the progeny as such deficiencies are unlikely to be rectified.

In recent years, more women are postponing childbearing to later ages, bringing the problems of reproductive aging into focus. Also, there is a surge in cases of ovarian disorders like Polycystic Ovarian Syndrome (PCOS) (Deswal et al., 2020) that affect oocyte quality (Patel and Carr, 2008). It therefore becomes imperative to study pathways involved in oocyte quality maintenance to delay reproductive aging. The current study emphasizes the crucial role of the interplay between cell-cycle proteins and nutrient signaling in shaping reproductive outcomes. Our findings uncover a previously unidentified role of somatic *cyclin D/cyd-1* in regulating reproductive fitness. Additionally, upon sensing cell cycle perturbations, activated DAF-16 destroyed somatic gonadal tissues, preventing the formation of low-quality oocytes. From an evolutionary standpoint, such mechanisms could be a strategy for the elimination of faulty progeny ensuring survival and propagation of healthy offspring in a species. In the future, it will be compelling to explore the mechanism through which cyclin D/CYD-1 and FOXO/DAF-16 impact such soma-germline signaling.

## Materials and Methods

### C. elegans strain maintenance

Unless otherwise mentioned, all the *C. elegans* strains were maintained and propagated at 20°C on *E. coli* OP50 using standard procedures (Stiernagle, 2006). The strains used in this study are:

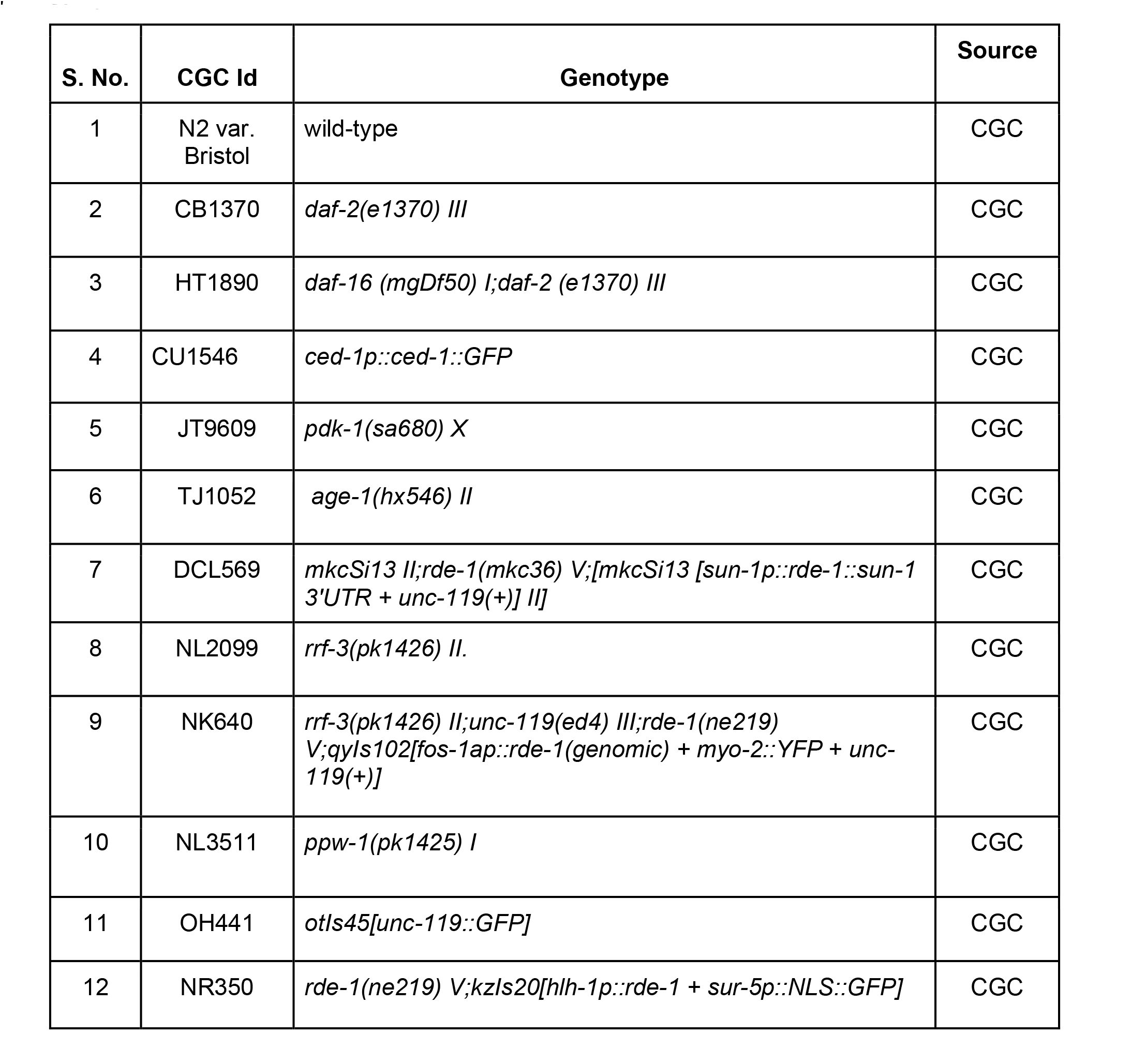
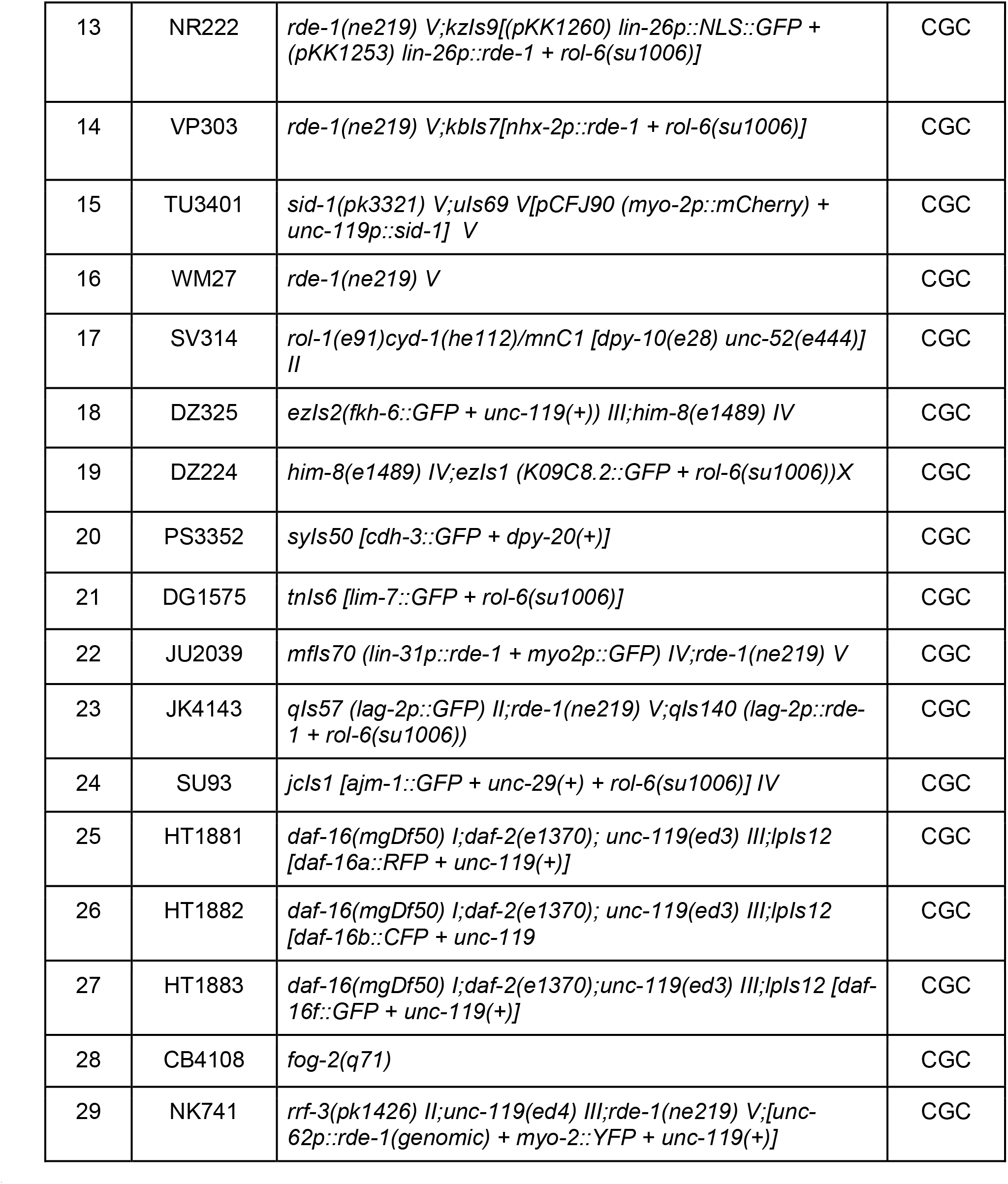

The strains listed below were generated in-house using standard cross-over techniques:

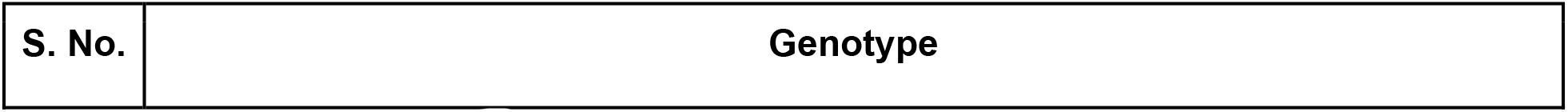
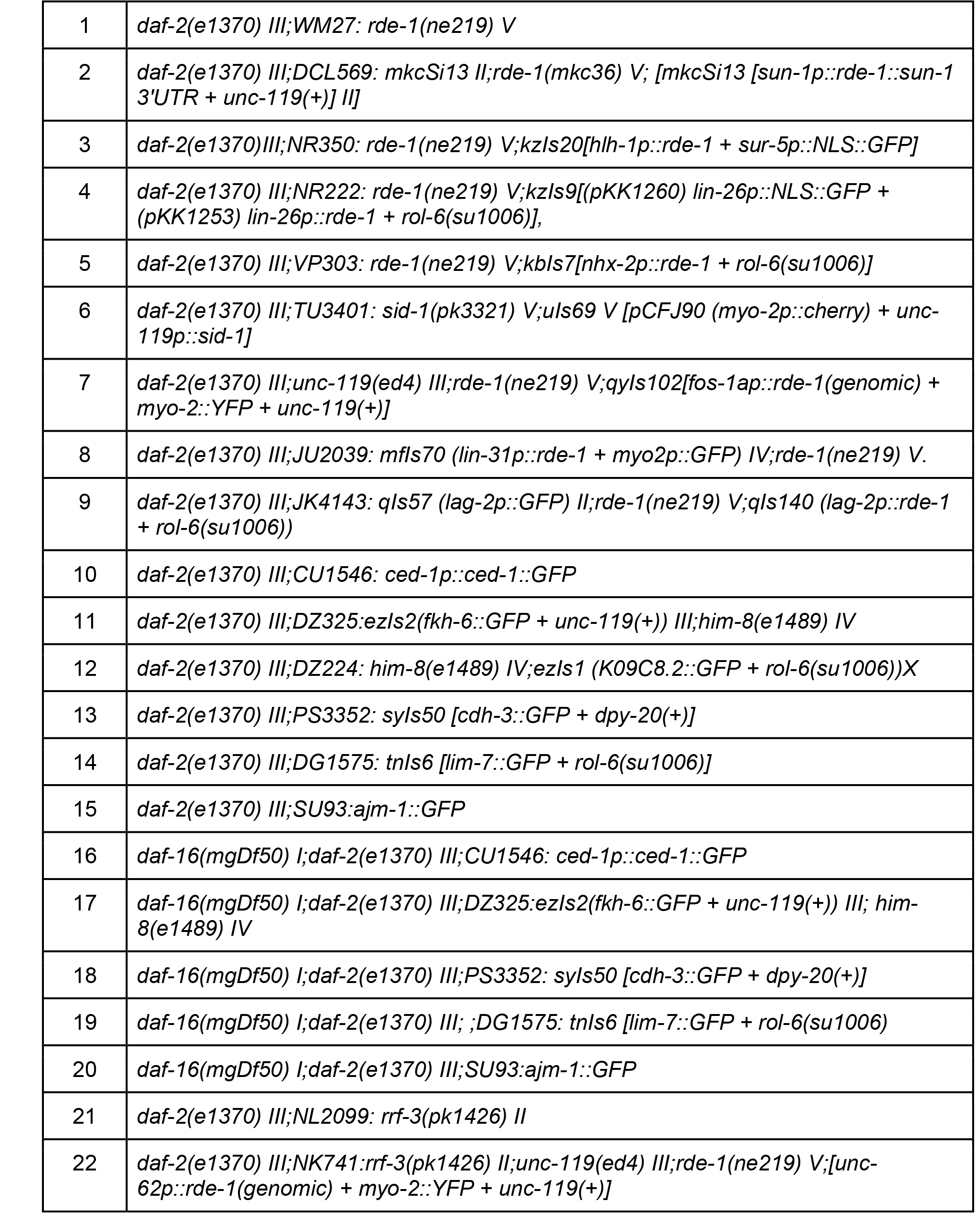

### Preparation of RNAi plates

RNAi plates were poured using an autoclaved nematode growth medium supplemented with 100 µg/ml ampicillin and 2 mM IPTG. Plates were allowed to dry at room temperature for 1 day. Bacterial culture harboring a particular RNAi construct was grown in Luria Bertani (LB) media supplemented with 100 µg/ml ampicillin and 12.5 µg/ml tetracycline, overnight at 37°C in a shaker incubator. Saturated primary cultures were re-inoculated the next day (1/50^th^ volume) in fresh LB media containing 100 µg/ml ampicillin and grown in a 37°C shaker until OD600 reached 0.5-0.6. The bacterial cells were pelleted down by centrifuging the culture at 3214 g for 5 minutes at 4°C and resuspended in 1/10^th^ volume of M9 buffer containing 100 µg/ml ampicillin and 1 mM IPTG. Around 300 µl of this suspension was seeded onto RNAi plates and left at room temperature for 2 days for drying, followed by storage at 4°C till further use.

### Hypochlorite treatment to obtain synchronize worm population

Gravid adult worms, initially grown on *E. coli* OP50 bacteria, were collected using M9 buffer in a 15 ml falcon tube. Worms were washed twice by first centrifuging at 652 g for 60 seconds followed by resuspension of the worm pellet in 1X M9 buffer. After the second wash, the worm pellet was resuspended in 3.5 ml of 1X M9 buffer and 0.5 ml 5N NaOH and 1 ml of 4% Sodium hypochlorite solution was added. The mixture was vortexed for 4-5 minutes until the entire worm body dissolved, leaving behind the eggs. The eggs were washed 5-6 times, by first centrifuging at 1258 g, the 1X M9 was aspirated out, followed by resuspension in fresh 1X M9 buffer to remove traces of bleach and alkali. After the final wash, eggs were kept in 15 ml falcons with ~ 10 ml of 1X M9 buffer and kept on rotation ~15 r.p.m for 17 hours to obtain L1 synchronized worms for all strains. The L1 worms were obtained by centrifugation at 805 g followed by resuspension in approximately 200-300 µl of M9 and added to respective RNAi plates.

### RNA isolation

Worms grown on respective RNAi were collected using 1X M9 buffer and washed thrice to remove bacteria. Trizol reagent (200 μl; Takara Bio, Kusatsu, Shiga, Japan) was added to the 50 μl worm pellet and subjected to three freeze-thaw cycles in liquid nitrogen with intermittent vortexing for 1 minute to break open the worm bodies. The samples were then frozen in liquid nitrogen and stored at −80 °C till further use. Later, 200 μl of Trizol was again added to the worm pellet and the sample was vigorously vortexed for 1 minute. To this, 200 μl of chloroform was added and the tube was gently inverted several times followed by 5 minutes of incubation at room temperature. The sample was then centrifuged at 12000 g for 15 minutes at 4°C. The RNA containing the upper aqueous phase was gently removed into a fresh tube without disturbing the bottom layer and interphase. To this aqueous solution, an equal volume of isopropanol was added and the reaction was allowed to sit for 10 minutes at room temperature followed by centrifugation at 12000 g for 15 minutes at 4°C. The supernatant was carefully discarded without disturbing the RNA-containing pellet. The pellet was washed using 1 ml 70% ethanol solution followed by centrifugation at 8000g for 10 minutes at 4°C. The RNA pellet was further dried at room temperature and later dissolved in autoclaved MilliQ water followed by incubation at 65°C for 10 minutes with intermittent tapping. The concentration of RNA was determined by measuring absorbance at 260 nm using a NanoDrop UV spectrophotometer (Thermo Scientific, Waltham, USA) and the quality was checked using denaturing formaldehyde-agarose gel.

### Gene expression analysis using quantitative real-time PCR (QRT-PCR)

First-strand cDNA synthesis was carried out using the Iscript cDNA synthesis kit (Biorad, Hercules, USA) following the manufacturer’s guidelines. The prepared cDNA was stored at −20°C. Gene expression levels were determined using the Brilliant III Ultra-Fast SYBR® Green QPCR master mix (Agilent, Santa Clara, USA) and Agilent AriaMx Real-Time PCR system (Agilent, Santa Clara, USA), according to manufacturer’s guidelines. The relative expression of each gene was determined by normalizing the data to actin expression levels. The list of primers is summarized in Table S4.

### Measurement of cell corpses using CED-1::GFP

The analysis of engulfed cell corpses was performed by utilizing transgenic worms expressing CED-1::GFP, a transmembrane protein found on surrounding sheath cells that are responsible for engulfing cell corpses. *ced-1p::ced-1::GFP(smIs34)* worms were bleached and their eggs were allowed to hatch in 1X M9 buffer for 17 hours to obtain L1 synchronized worms. Approximately 100 L1 worms were placed onto control or *cyd-1* RNAi in triplicates. On the day-1 of adulthood, worms were imaged in Z-stack using 488 nm laser to excite GFP on LSM980 confocal microscope (Carl Zeiss, Oberkochen, Germany). The images were converted into a maximum intensity projection (MIP). The number of cell corpses per gonadal arm was quantified manually.

### DAPI staining

Worms were cultured on specific RNAi plates from the L1 stage onward. On day-1 of adulthood, worms were collected in a 1.5 ml Eppendorf tube with 1X M9 buffer and allowed to settle. Using a glass Pasteur pipette, the 1X M9 buffer was carefully removed, leaving approximately 100 µl of worm suspension. Subsequently, 1 ml of ice-cold 100% methanol was added to the worm pellet, and the mixture was incubated at −20°C for 30 minutes. The methanol- fixed worm pellet was then placed on a glass slide, and after the methanol had evaporated, Fluoroshield with DAPI (from Invitrogen, Carlsbad, USA) was applied for staining. A coverslip was carefully placed on top (avoiding bubbles) and sealed with transparent nail paint.

To stain dissected gonads, worms were positioned on a glass slide in 1X M9, and the gonads were obtained by carefully cutting the pharynx or tail end of the worm using a sharp 25G needle. After this 500 µl of chilled 100% methanol was added to the slide and allowed to evaporate. Subsequently, Fluoroshield with DAPI (Invitrogen, Carlsbad, USA) was added for staining and a coverslip was carefully placed on top (avoiding bubbles) and sealed with transparent nail paint. Finally, the images were captured using a 405 nm laser excitation DAPI on LSM980 confocal microscope (Carl Zeiss in Oberkochen, Germany).

### Brood size, reproductive span, and egg hatching

#### Brood size

Worms were grown on control or *cyd-1* RNAi from L1 onwards and upon reaching the young adult stage, five worms were picked onto fresh RNAi plates, in triplicates, and allowed to lay eggs for 24 hours. The worms were then transferred to fresh plates every day until worms ceased to lay eggs, and the eggs laid on the previous day’s plate were counted. These plates were again counted after 48 hours to document the number of hatched worms. The total number of hatched progenies per worm is defined as brood size.

#### Reproductive span

Every day, individual L4 hermaphrodites were relocated to fresh RNAi plates until they laid no eggs for a minimum of two consecutive days. The final day when they were capable of producing viable offspring was recorded as their reproductive span. Each experiment included a minimum of 15 worms per condition. The log-rank (Mantel-Cox) method was employed to assess the null hypothesis.

#### Egg hatching

The quality of eggs was assessed by determining the percentage of hatched eggs. Almost 50 adult worms per condition were allowed to lay eggs for an hour. The mother worms were then sacrificed and the number of eggs on respective plates were counted. After 48 hours the number of hatched progenies were counted and the percentage of hatched eggs were calculated for each condition.

#### Analysis of Oocyte Morphology

WT or *rrf-3(pk1426)* worms were grown from L1 onwards on control or *cyd-1* RNAi. Microscopic images of oocytes on the third day of adulthood were captured using differential interference contrast (DIC) settings (Carl Zeiss, Oberkochen, Germany). These oocytes were classified into three groups based on various characteristics such as cavities, shape, size, and organization. Depending on the severity of the observed features, oocytes were classified as either normal, mild, or severe. Oocytes without cavities, small size, or disorganization were labeled as normal. Oocytes exhibiting the presence of either two small oocytes, two cavities, or two disorganized oocytes were categorized as having a mild phenotype. Those with more than two cavities, small oocytes, or misshapen oocytes were considered to have a severe phenotype.

#### Quantification of fertile worms

Approximately 100 synchronized L1 worms were placed onto different RNAi plates, in triplicates. On the day-1 adult stage, bright-field images were captured (Carl Zeiss, Oberkochen, Germany). Worms carrying over five eggs in their uterus were categorized as fertile.

#### Germ cell count

Around 100 L1-staged worms were poured onto control or *cyd-1* RNAi plates in triplicate. Utilizing a 405 nm laser to excite DAPI in a confocal microscope (Carl Zeiss, Oberkochen, Germany), images of DAPI stained day-1 adult germline were acquired. Z stacked images (0.13 µM sections between the starting plane and end plane) of germline were obtained followed by Maximum Intensity projection (MIP) of each z-stack in the XY plane to create a superimposed image. Quantification of germ cells per gonadal arm was done using Fiji (National Institute of Health) in different regions based on their morphology, namely the Mitotic Zone (spanning from the distal tip to the transition zone), the Transition zone (extending from the beginning to the end of crescent-shaped nuclei), and the pachytene zone (from the end of the transition zone to the turn region).

#### Sperm count

DAPI stained early day-1, day-2 or day-3 adult gonads were imaged using 405 nm laser to excite DAPI on a confocal microscope (Carl Zeiss, Oberkochen, Germany). Z stacked images (0.13 µM sections between the starting plane and end plane) of germline were obtained followed by Maximum Intensity projection (MIP) of each z-stack in the XY plane to create a superimposed image. Sperms per gonadal arm were quantified using Fiji (National Institute of Health).

#### Scoring of Endomitotic oocyte (emo)

Approximately 50 L1-staged worms were placed onto control or *cyd-1* RNAi plates in triplicate. Day-3 adult worm gonads were imaged using differential interference contrast (DIC) settings in a microscope (Carl Zeiss, Oberkochen, Germany). Endomitotic oocytes in the uterus were identified by following features: a lack of distinct nucleus, nuclei may appear disorganized, lobulated, or enlarged (McCarter et al., 1997). Alternatively, day-3 adults were DAPI stained and subsequently imaged in a Z-stack using 405 nm laser to excite DAPI on an LSM980 confocal microscope (Carl Zeiss, Oberkochen, Germany). Nuclei that appeared enlarged and exhibited dense DAPI staining (Iwasaki et al. in 1996), were identified as endomitotic oocytes (emo). For the purpose of scoring, the images were converted into a maximum intensity projection (MIP).

#### unc-119::gfp positive emo

Approximately 50 L1-staged *unc-119::gfp* (neuronal fate marker) transgenic worms were placed onto control or *cyd-1* RNAi plates in triplicates. Day-3 adult worm gonads were imaged using 488 nm laser to excite GFP as well as differential interference contrast (DIC) settings in a fluorescence microscope (Carl Zeiss, Oberkochen, Germany). Endomitotic oocytes in the uterus were identified by following features: a lack of distinct nucleus, nuclei may appear disorganized, lobulated or enlarged (McCarter et al., 1997). The number of worms having endomitotic oocytes as well as that expressed *unc-119::gfp* in the emo were counted. Percentage of worms expressing GFP in emo was calculated for each condition.

#### Imaging of transgenic reporter strains

Synchronized L1 populations of transgenic reporter strains were grown on respective RNAi plates. For imaging, the worms were mounted on agar pads with 20 mM sodium azide and a coverslip was placed on the paralyzed worms. The coverslip was secured with a transparent nail paint. All images were acquired in a Z-stack using 488 nm laser to excite GFP on LSM980 confocal microscope (Carl Zeiss, Oberkochen, Germany). The images were converted into a maximum intensity projection (MIP). To qualitatively quantify *lim-7p::gfp* (sheath cell marker) and *fkh-6::gfp* (hermaphrodite spermatheca) expression, the day-1 or day-3 adult gonads were scored as having normal expression, reduced expression (less intensity than normal) or missing (no GFP expression). In the case of *cdh-3::gfp* (a marker for vulval cells), we used late-L3 staged worms and scored their gonads based on whether they displayed normal expression, abnormal expression (distorted pattern), or missing (no GFP expression). Similar categorization was utilized to assess the *ajm-1::gfp* (adherens junction marker) expression in early day-1 adult gonads.

#### Phalloidin staining and sheath cell structure

Worms were grown on control and *cyd-1* RNAi L1 onwards. Day-1 or day-3 adult worms were collected in a 1.5 ml Eppendorf tube with 1X M9 buffer and allowed to settle. Using a glass Pasteur pipette, excess 1X M9 buffer was carefully removed, and worms were transferred to a glass slide. Using a 25G needle, worms were excised at the pharynx or near tail region, causing gonad arms to extrude from the body. The dissected gonad arms were transferred to 5 ml glass tubes and fixed in 4% formaldehyde for 20 minutes at room temperature and then washed thrice with 0.1% 1X PBST. After washing, fixed gonad arms were transferred to 1 ml glass tubes and stained with Oregon Green™ 488 Phalloidin (Molecular Probes, Invitrogen, Life Technologies, Grand Island, NY, USA) at a final concentration of 0.4 units/ml and placed in the dark at 4°C overnight followed by 30 minutes incubation at RT. Gonad arms were then washed twice with PBST, and extra liquid was removed and gonad arms were transferred onto slides with a drop of Fluoroshield with DAPI (Invitrogen, Carlsbad, USA). Coverslip was placed on the mounted worms and sealed using nail-paint. Phalloidin marks the actin structure. To analyze sheath cell structure gonads were imaged in a Z-stack using 488 nm laser to excite GFP on LSM980 confocal microscope (Carl Zeiss, Oberkochen, Germany). The images were converted into a maximum intensity projection (MIP). Sheath cell structure was scored as normal or mild defect (slight clustering or loosening of actin filaments) or severe defect (highly clustered or loosened actin filaments). To examine vulval morphology, the images were captured with a focus on the vulva- uterine region.

#### Vulva morphology

Worms were grown from L1 onwards on control or *cyd-1* RNAi. Microscopic images of vulva on the early day-1 of adulthood were captured using differential interference contrast (DIC) settings on a microscope (Carl Zeiss, Oberkochen, Germany). These vulvas were classified as normal or abnormal (protruding vulva/vulvaless) based on its structure and organization.

#### RNA sequencing

Synchronized late-L4 worms grown on control or *cyd-1* RNAi were collected using 1X M9 buffer, after washing it thrice to remove bacteria. Total RNA was isolated from these worm pellets using the Trizol method. The concentration of RNA was determined by measuring absorbance at 260 nm using a NanoDrop UV spectrophotometer (Thermo Scientific, Waltham, USA) and RNA quality was checked using RNA 6000 NanoAssay chip on a Bioanalyzer 2100 machine (Agilent Technologies, Santa Clara, USA). RNA above RNA integrity number = 9 was included in the study. An 1.5 μg aliquot of RNA samples was used for polyA mRNA isolation from total RNA using NEBNext® Poly(A) mRNA Magnetic Isolation Module (Catalog no-E7490L) according to the maufacturer’s protocol. The NEBNext® Ultra™ II Directional RNA Library Prep kit (Catalog no- E7760L) was used to construct libraries according to the manufacturer’s instructions. NovaSeq6000 NGS platform (Illumina Inc., San Diego, California, USA) with 2X150 paired end chemistry was used to generate 30 million paired end reads corresponding to 9 Gb data per sample. The RNA sequencing data are available at https://www.ebi.ac.uk/biostudies/arrayexpress with accession number E-MTAB-13172.

#### RNA-seq Analysis

Sequencing reads were subjected to quality control using the Fastp kit (Chen et al., 2018). The quality of reads was assessed and visualized by FASTQC (https://www.bioinformatics.babraham.ac.uk/projects/fastqc). Adapter sequence was removed by Trimmomatic tool (Bolger et al., 2014). For each of our samples, reads were aligned to the WBcel235 cell genome using STAR-2.7.10b (Dobin and Gingeras, 2015) with an average 94.4% alignment rate. Gene counts were obtained with the quantMode GeneCounts option. Differential Gene expression analysis was performed using DeSeq2 (Love et al., 2014) package in R. Differentially expressed genes were defined as those with padj values below 0.06. Genes with a cut-off of log2(fold change) > 0.6 and log2(fold change) < −0.6 were considered as upregulated and downregulated genes, respectively. For downstream analysis, the function variance stabilizing transformations (VST) in the DeSeq2 package was implemented. Wormbase GeneIDs were mapped to their gene symbols by AnnotationaDbi and *C. elegans* database (org.Ce.eg.db) packages in R. Counts function was used to normalize the gene counts. Gene Set Enrichment analysis was performed using gseGO function of clusterProfiler v4.6.2 (Wu et al., 2021) in R. Volcano plots and dot plots were plotted with ggplot2 (https://ggplot2.tidyverse.org) in R. For volcano plots, differentially expressed genes (padj <0.1 and |log2(fold change)|>0.6) were indicated in green and purple colors. Dot plots were made with Enriched GO terms on y-axis and Normalized Enrichment Score (NES) on x-axis and size and color as padj value.

#### Statistical tests

All statistical tests were performed utilizing the built-in functions of GraphPad Prism 10.1.0. In instances where we compared two conditions, we applied a Student’s *t*-test with Welch’s correction, making no assumptions about consistent standard deviations. When comparing multiple conditions, we employed a Two-way ANOVA with Tukey’s multiple comparison test.

## Acknowledgements

We thank Ranjisha KR and Nikhita Anand for their help with experiments, and all members of Molecular Aging Laboratory for their support. The SU93 strain *jcIs1[AJM-1::GFP + unc-29(+) + rol-6(su1006)] IV* was kindly provided by Dr. K. Subramaniam and we also thank him for his valuable suggestions.

## Data availability statement

The data reported in this manuscript is available in the Source data file that accompanies the manuscript as Table S1.

The RNA seq reads have been submitted https://www.ebi.ac.uk/biostudies/arrayexpress with the accession number E-MTAB-13172.

## Author Contributions

UR, GCS, AM conceptualized the project. UR, GCS, AC, DC performed the experiments and analyzed data. MR performed bioinformatics analysis under the supervision of AGA. UR, GCS, AM wrote the manuscript. AM supervised the project and acquired funding.

## Classification

Biological Sciences: Developmental Biology.

