## Supplementary Figure for "A non-canonical role of somatic CYCLIN D/CYD-1 in oogenesis and reproductive aging, dependent on the FOXO/DAF-16 activation state"

**Author Contributions:** UR, GCS, AM conceptualized the project. UR, GCS, AC, DC performed the experiments and analyzed data. MR performed bioinformatics analysis under the supervision of AGA. UR, GCS, AM wrote the manuscript. AM supervised the project and acquired funding.

**Competing Interest Statement:** The authors disclose that there is no competing interest.

**Classification:** Biological Sciences: Developmental Biology.

Keywords: Cyclin D/CYD-1, FOXO/DAF-16, insulin signaling, pachytene arrest, germ line, somatic gonad.

This PDF file includes: Figure S1-6 along with their legends.

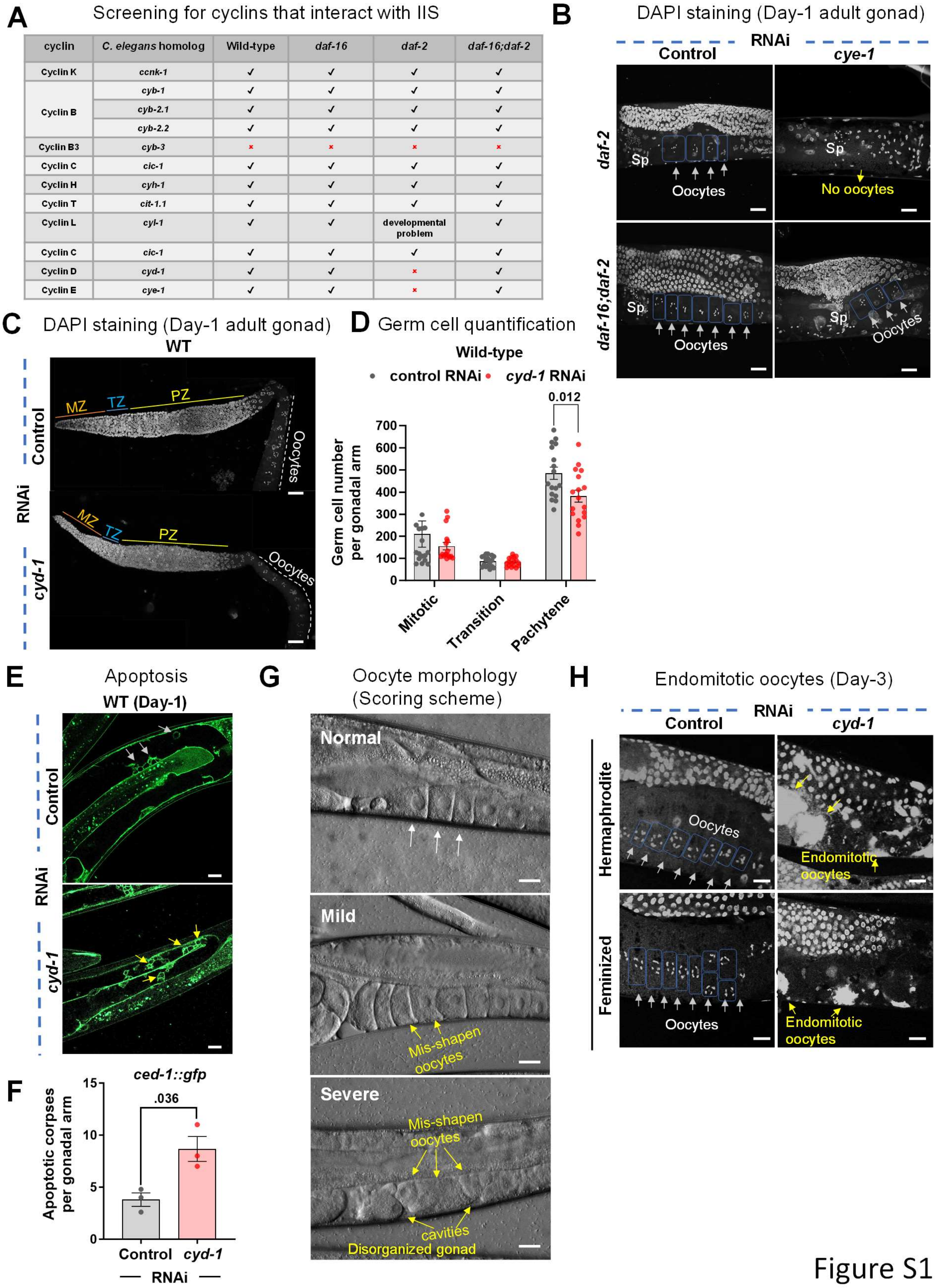


| cyclin | *C. elegans* homolog | Wild-type | *daf-16* | *daf-2* | *daf-16;daf-2* |
|---|---|---|---|---|---|
| Cyclin K | *ccnk-1* | ✓ | ✓ | ✓ | ✓ |
| Cyclin B | *cyb-1* | ✓ | ✓ | ✓ | ✓ |
| | *cyb-2.1* | ✓ | ✓ | ✓ | ✓ |
| | *cyb-2.2* | ✓ | ✓ | ✓ | ✓ |
| Cyclin B3 | *cyb-3* | × | × | × | × |
| Cyclin C | *cic-1* | ✓ | ✓ | ✓ | ✓ |
| Cyclin H | *cyh-1* | ✓ | ✓ | ✓ | ✓ |
| Cyclin T | *cit-1.1* | ✓ | ✓ | ✓ | ✓ |
| Cyclin L | *cyl-1* | ✓ | ✓ | developmental problem | ✓ |
| Cyclin C | *cic-1* | ✓ | ✓ | ✓ | ✓ |
| Cyclin D | *cyd-1* | ✓ | ✓ | × | ✓ |
| Cyclin E | *cye-1* | ✓ | ✓ | × | ✓ |

Figure S1

**Supplementary Figure 1**

**(A)** RNAi screen for cyclins that may interact with the IIS. Twelve cyclin genes were knocked down in WT, *daf-16(mgdf50)*, *daf-2(e1370)* and *daf-16(mgdf50);daf-2(e1370)* using RNAi. The tick mark indicates fertile worms while the cross mark indicates sterility when grown on respective RNAi from L1 onwards.

**(B)** Representative fluorescent images of DAPI-stained germ line of *daf-2(e1370)* and *daf-16(mgdf50);daf-2(e1370)* worms grown on control or *cye-1* RNAi. Oocytes are boxed for clarity. White arrows point towards oocytes while yellow shows the absence of oocytes. Sp denotes sperms.

**(C,D)** Representative images of DAPI-stained dissected gonadal arm of WT worms (day-1 adult) grown on control or *cyd-1* RNAi. Mitotic zone (MT), transition Zone (TZ), and pachytene zone (PZ) germ cells are marked with a solid line and white dashed line mark oocytes (C). Quantitation of germ cells in each zone (D). n = 17 gonads for each condition. Each point represents the number of germ cells in the respective zones. Unpaired *t*-test with Welch's correction.

**(E,F)** Representative images showing apoptotic cells(marked by arrows) in the gonadal arm of *ced-1::gfp* worms (day-1 adult) grown on control or *cyd-1* RNAi (E). Quantification for apoptotic cells per gonadal arm (F) The average of three biological replicates is shown (n ≥ 15 for each replicate). Unpaired *t*-test with Welch's correction.

**(G)** Oocyte quality score based on morphology. The quality was categorized as normal, or mild or severe based on the presence of cavities, shape and organization of oocytes. Normal = No cavities/not misshapen/not disorganized, mild = either with cavities/ are misshapen/disorganized (≤ 2 instances per worm), severe= either with cavities/ are misshapen/disorganized (≥ 3 instances per worm).

**(H)** Representative DAPI-stained gonads of WT hermaphrodites (day-3 adult) and *fog-2(q71)* females grown on control or *cyd-1* RNAi at 25℃. Oocytes are boxed for clarity. White arrows point towards oocytes while yellow arrows point towards endomitotic oocytes (emos).

Scale bars:20 μm. Error bars are s.e.m. Experiments were performed at 20°C except S1H which was performed at 25°C. Source data are provided in Table S1.

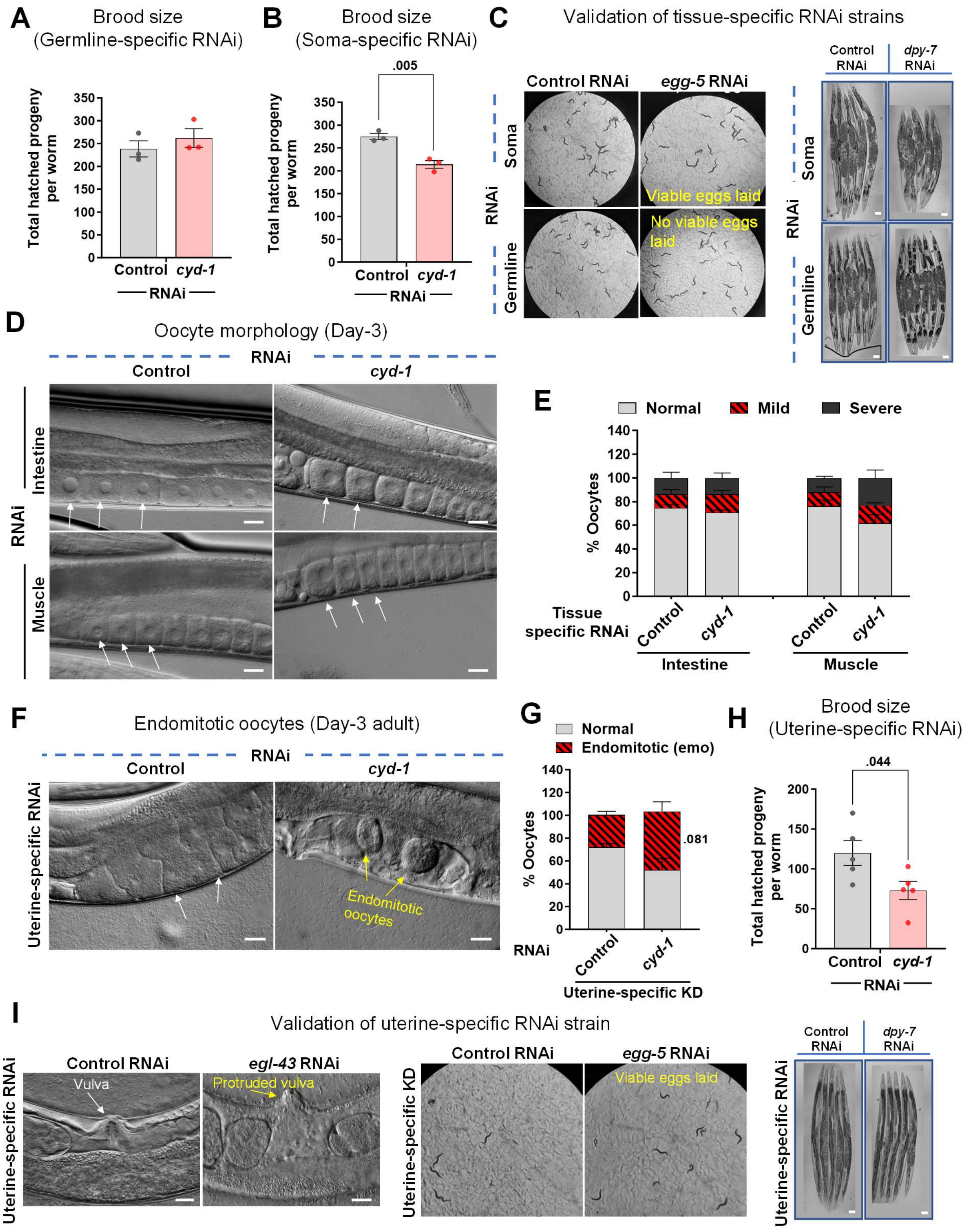
A
Brood size
(Germline-specific RNAi)
Total hatched progeny per worm
Control
cyd-1
RNAi
B
Brood size
(Soma-specific RNAi)
.005
Total hatched progeny per worm
Control
cyd-1
RNAi
C
Validation of tissue-specific RNAi strains
Control RNAi
egg-5 RNAi
Soma
Germline
RNAi
Viable eggs laid
No viable eggs laid
Control RNAi
dpy-7 RNAi
D
Oocyte morphology (Day-3)
RNAi
Control
cyd-1
Intestine
Muscle
E
Normal
Mild
Severe
% Oocytes
Tissue specific RNAi
Control
cyd-1
Intestine
Muscle
F
Endomitotic oocytes (Day-3 adult)
RNAi
Control
cyd-1
Uterine-specific RNAi
Endomitotic oocytes
G
Normal
Endomitotic (emo)
% Oocytes
.081
RNAi
Control
cyd-1
Uterine-specific KD
H
Brood size
(Uterine-specific RNAi)
.044
Total hatched progeny per worm
Control
cyd-1
RNAi
I
Validation of uterine-specific RNAi strain
Control RNAi
egl-43 RNAi
Vulva
Protruded vulva
Uterine-specific RNAi
Uterine-specific KD
Control RNAi
egg-5 RNAi
Viable eggs laid
Control RNAi
dpy-7 RNAi

Figure S2

**Supplementary Figure 2.**

**(A)** The total number of hatched progenies in *rde-1(mkc36);sun-1p::rde-1* worms (germline-specific RNAi) grown on control or *cyd-1* RNAi. Average of three biological repeats. Unpaired *t*-test with Welch's correction.

**(B)** The total number of hatched progenies in *ppw-1(pk1425)* worms (soma-specific RNAi) grown on control or *cyd-1* RNAi. Average of three biological repeats. Unpaired *t*-test with Welch's correction.

**(C)** Representative images of *ppw-1(pk1425)* (soma-specific RNAi), or *rde-1(mkc36);sun-1p::rde-1* (germline-specific RNAi) worms (day-1 adult) grown on control, *egg-5* or *dpy-7* RNAi. KD of *egg-5* led to non-viable progeny in the germline-specific RNAi strain but not in the soma-specific RNAi strain, while KD of *dpy-7* led to dumpy phenotype in the soma-specific RNAi strain but not in the germline-specific RNAi strain. Scale bar 100 µm.

**(D,E)** DIC images showing oocyte morphology of *rrf-3(pk1426);rde-1(ne219);nhx-2p::rde-1* (intestine-specific RNAi) or *rrf-3(pk1426);rde-1(ne219);hlh-1p::rde-1* (muscle-specific RNAi) worms (day-3 adult) grown on control or *cyd-1* RNAi. White arrows mark oocytes. Scale bar 20 µm (D). Oocyte quality score (based on morphology as shown in Figure S1G) (E). Average of three biological replicates (n ≥ 25 for each replicate). Unpaired *t*-test with Welch's correction.

**(F,G)** DIC images showing the presence of unfertilized oocytes/endomitotic oocytes (emos) in the uterus of *rrf-3(pk1426);rde-1(ne219);fos-1ap::rde-1(genomic)* (uterine tissue-specific RNAi) worms (day-3 adult) grown on control or *cyd-1* RNAi. White arrows mark unfertilized oocytes in the uterus while yellow arrows mark endomitotic oocytes (emos). Scale bar 20 µm (F). Quantification for endomitotic oocytes (G) Average of three biological replicates (n ≥ 25 for each experiment). Unpaired *t*-test with Welch's correction.

**(H)** The total number of hatched progenies in *rrf-3(pk1426);rde-1(ne219);fos-1ap::rde-1(genomic)* (uterine tissue-specific RNAi) worms grown on control or *cyd-1* RNAi. Average of four biological repeats. Unpaired *t*-test with Welch's correction.

**(I)** Representative images of *rrf-3(pk1426);rde-1(ne219);fos-1ap::rde-1* (uterine tissue-specific RNAi) worms (day-1 adult) grown on *egl-43, egg-5 or dpy-7* RNAi. KD of *egl-43* led to protruded vulva, while *egg-5* KD resulted in viable progeny and *dpy-7* KD led to non-dumpy progeny. Scale bar 100 µm.

Error bars are s.e.m. Experiments were performed at 20°C. Source data are provided in Table S1.

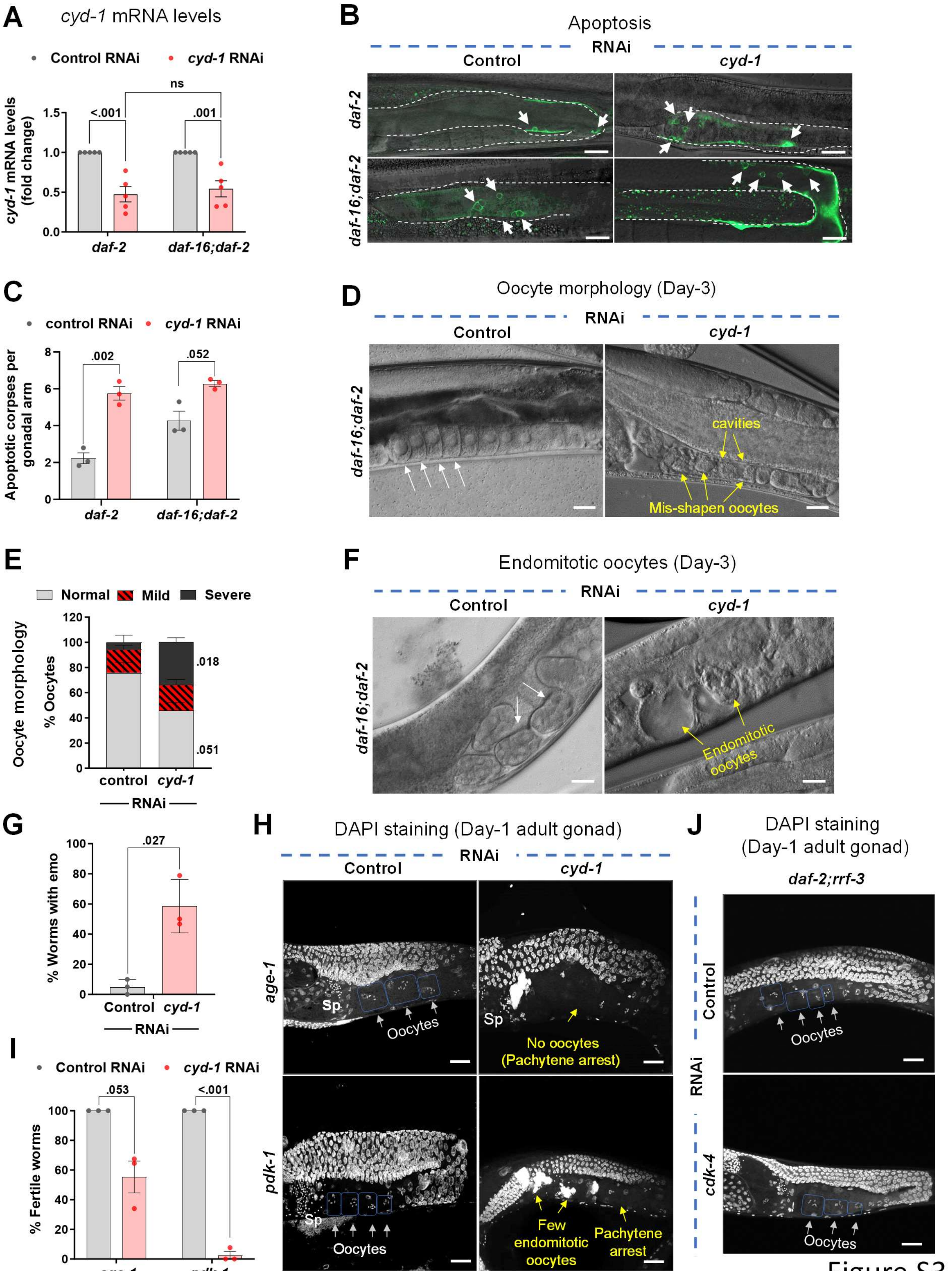
A
cyd-1 mRNA levels
Control RNAi
cyd-1 RNAi
cyd-1 mRNA levels (fold change)
ns
<.001
.001
1.5
1.0
0.5
0.0
daf-2
daf-16;daf-2
B
Apoptosis
RNAi
Control
cyd-1
daf-2
daf-16;daf-2
C
control RNAi
cyd-1 RNAi
Apoptotic corpses per gonadal arm
.002
.052
8
6
4
2
0
daf-2
daf-16;daf-2
D
Oocyte morphology (Day-3)
RNAi
Control
cyd-1
daf-16;daf-2
cavities
Mis-shapen oocytes
E
Normal
Mild
Severe
Oocyte morphology
% Oocytes
120
100
80
60
40
20
0
.018
.051
control
cyd-1
RNAi
F
Endomitotic oocytes (Day-3)
RNAi
Control
cyd-1
daf-16;daf-2
Endomitotic oocytes
G
% Worms with emo
.027
100
80
60
40
20
0
Control
cyd-1
RNAi
H
DAPI staining (Day-1 adult gonad)
RNAi
Control
cyd-1
age-1
Sp
Oocytes
No oocytes (Pachytene arrest)
pdk-1
Few endomitotic oocytes
Pachytene arrest
I
Control RNAi
cyd-1 RNAi
% Fertile worms
.053
<.001
100
80
60
40
20
0
age-1
pdk-1
J
DAPI staining (Day-1 adult gonad)
daf-2;rrf-3
Control
RNAi
cdk-4
Oocytes

Figure S3

**Supplementary Figure 3**

**(A)** RT-PCR analysis showing knockdown efficiency of *cyd-1* RNAi in *daf-2(e1370)* and *daf-16(mgdf50);daf-2(e1370)*. Expression levels were normalized to *actin*. Averages of 5 biological replicates are shown. Two-way ANOVA-Tukey's multiple comparisons test.

**(B,C)** Representative fluorescent and DIC merged images showing apoptotic cells (arrows) in the gonadal arm of *daf-2(e1370);ced-1::gfp and daf-16(mgdf50);daf-2(e1370);ced-1::gfp* (day-1 adult) worms grown on control or *cyd-1* RNAi. Arrows mark apoptotic corpses. (B). Quantification for apoptotic corpses per gonadal arm (C). An average of three biological replicates are shown (n ≥ 17 for each replicate). Unpaired *t*-test with Welch's correction.

**(D,E)** DIC images showing oocyte morphology of *daf-16(mgdf50);daf-2(e1370)* (day-3 adult) worms grown on control or *cyd-1* RNAi. White arrows mark normal oocytes whereas yellow arrows mark oocytes with cavities or those that are misshapen or disorganized, indicative of poor quality (D). Oocyte quality score (based on morphology as shown in Figure S1G. ) (E). Average of three biological replicates (n ≥ 25 for each replicate). Unpaired *t*-test with Welch's correction.

**(F,G)** DIC images showing eggs or endomitotic oocytes (emos) in *daf-16(mgdf50);daf-2(e1370)* (day-3 adult) worms grown on control or *cyd-1* RNAi. White arrows mark normal eggs in the uterus whereas yellow arrows mark emos (F). Quantification for endomitotic oocytes (G). Average of three biological replicates (n ≥ 20 for each replicate). Unpaired *t*-test with Welch's correction.

**(H,I)** Representative fluorescent images of DAPI stained gonads of *age-1(hx546)* and *pdk-1(sa680)* (day-1 adult) worms grown on control or *cyd-1* RNAi. Oocytes are boxed for clarity. White arrows point towards oocytes while yellow shows the absence of oocytes. Sp denotes sperms (H). The percentage of fertile worms (I). Average of three biological replicates (n ≥ 30 for each replicate). Unpaired *t*-test with Welch's correction.

**(J)** Representative DAPI-stained gonads of *rrf-3(pk1426);daf-2(e1370)* worms (day-1 adult) grown on control or *cdk-4* RNAi. Oocytes are boxed for clarity. White arrows point towards oocytes. Scale bar 20 µm.

Scale bars:20 µm. Error bars are s.e.m. Experiments were performed at 20°C. Source data are provided in Table S1.

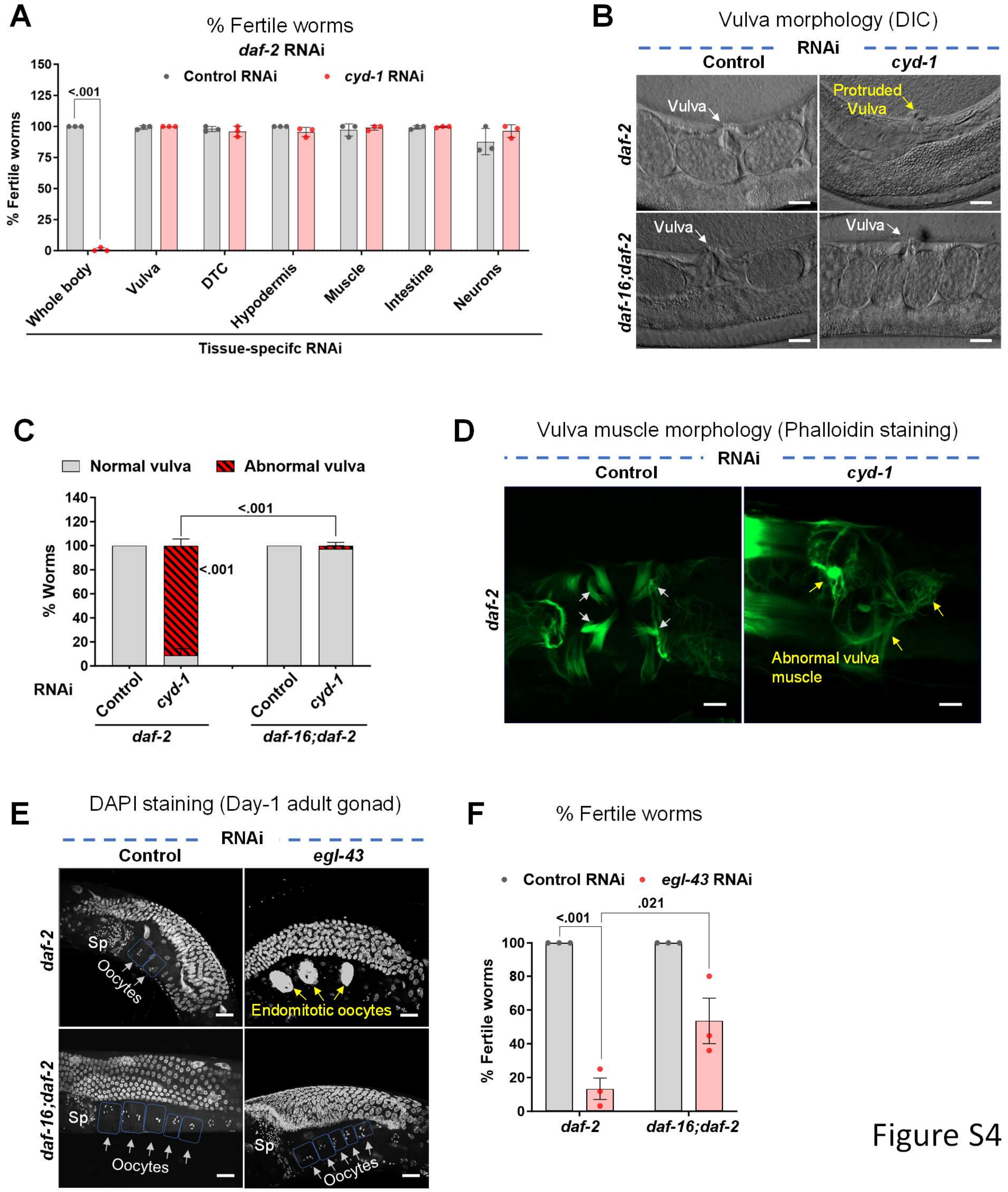
A
% Fertile worms
daf-2 RNAi
Control RNAi
cyd-1 RNAi
% Fertile worms
<.001
Whole body
Vulva
DTC
Hypodermis
Muscle
Intestine
Neurons
Tissue-specifc RNAi
B
Vulva morphology (DIC)
RNAi
Control
cyd-1
daf-2
daf-16;daf-2
Vulva
Protruded Vulva
C
Normal vulva
Abnormal vulva
% Worms
<.001
<.001
RNAi
Control
cyd-1
daf-2
daf-16;daf-2
D
Vulva muscle morphology (Phalloidin staining)
RNAi
Control
cyd-1
daf-2
Abnormal vulva muscle
E
DAPI staining (Day-1 adult gonad)
RNAi
Control
egl-43
daf-2
daf-16;daf-2
Sp
Oocytes
Endomitotic oocytes
F
% Fertile worms
Control RNAi
egl-43 RNAi
<.001
.021
% Fertile worms
daf-2
daf-16;daf-2

Figure S4

**Supplementary Figure 4.**

**(A)** The percentage of fertile worms upon tissue-specific *cyd-1* KD in *daf-2(e1370).* Average of three biological replicates (n ≥ 25 per condition for each experiment). Unpaired *t*-test with Welch's correction.

**(B,C)** DIC images of gonads of *daf-2(e1370)* and *daf-16(mgdf50);daf-2(e1370)* worms (day-1 adult) on control or *cyd-1* RNAi. White arrows point towards normal vulva while yellow points towards abnormal vulva (protruded). Scale bar 100 μm (B). Quantification of normal and abnormal vulva (C). Average of three biological replicates (n ≥ 30 for each experiment). Two-way ANOVA-Tukey's multiple comparisons test.

**(D)** Representative fluorescent images of phalloidin-stained (that marks the F-actin) gonads of *daf-2(e1370)* worms (day-1 adult) grown on control or *cyd-1* RNAi. White arrows show normal vulva muscle structure yellow arrows show defective vulva muscles. Scale bar 10 μm.

**(E,F)** Representative fluorescent images of DAPI-stained gonads of *daf-2(e1370)* and *daf-16(mgdf50);daf-2(e1370)* (day-1 adult) worms grown on control or *egl-43* RNAi. Oocytes are boxed for clarity. White arrows point towards oocytes while yellow shows endomitotic oocytes. Sp denotes sperms. Scale bar 20 μm (E). The percentage of fertile worms (F). Average of three biological replicates (n ≥ 25 per condition for each experiment). Two-way ANOVA-Tukey's multiple comparisons test.

Error bars are s.e.m. Experiments were performed at 20°C. Source data are provided in Table S1.

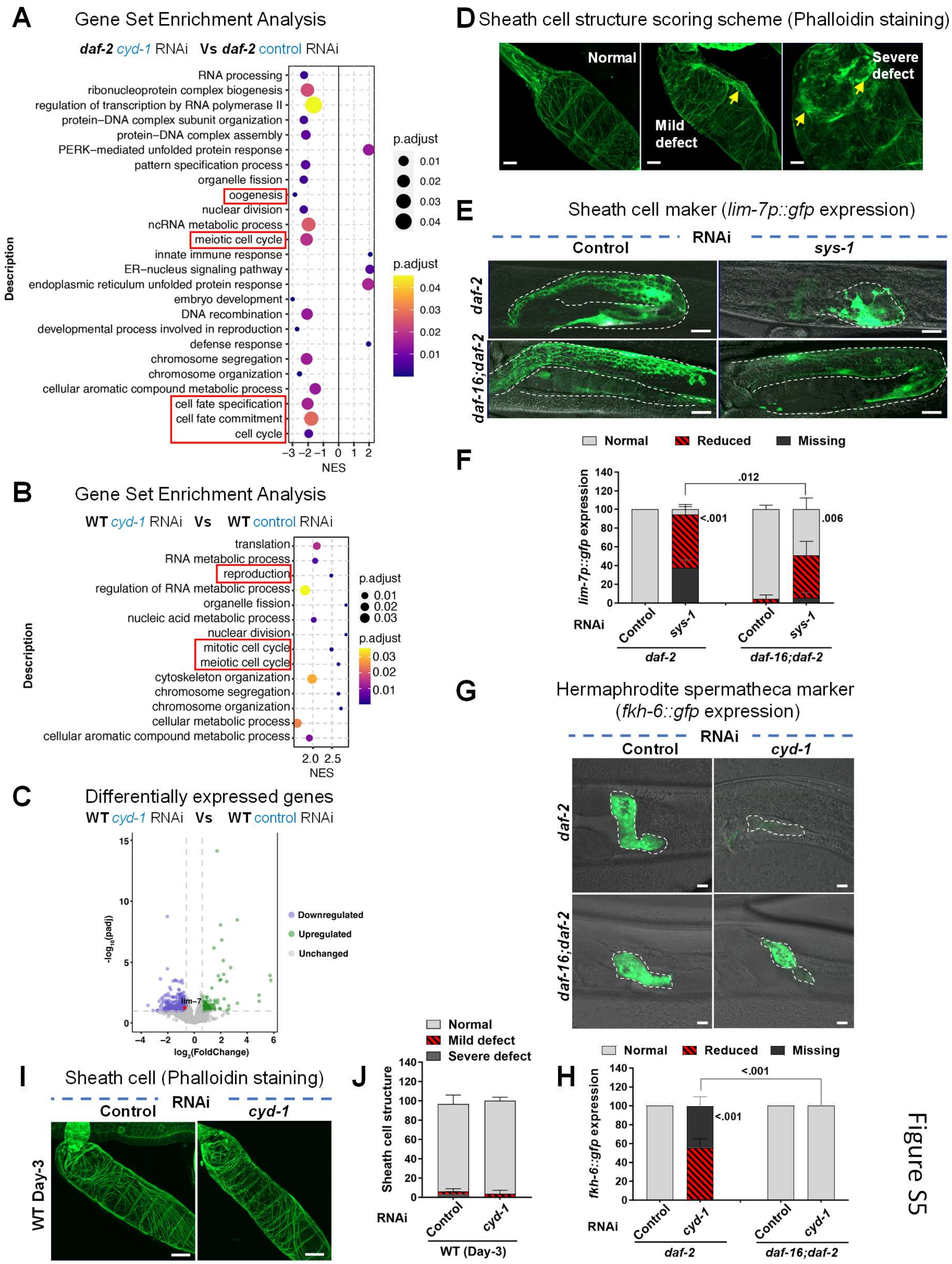
A
Gene Set Enrichment Analysis
daf-2 cyd-1 RNAi Vs daf-2 control RNAi
RNA processing
ribonucleoprotein complex biogenesis
regulation of transcription by RNA polymerase II
protein–DNA complex subunit organization
protein–DNA complex assembly
PERK–mediated unfolded protein response
pattern specification process
organelle fission
oogenesis
nuclear division
ncRNA metabolic process
meiotic cell cycle
innate immune response
ER–nucleus signaling pathway
endoplasmic reticulum unfolded protein response
embryo development
DNA recombination
developmental process involved in reproduction
defense response
chromosome segregation
chromosome organization
cellular aromatic compound metabolic process
cell fate specification
cell fate commitment
cell cycle
Description
NES
p.adjust
B
Gene Set Enrichment Analysis
WT cyd-1 RNAi Vs WT control RNAi
translation
RNA metabolic process
reproduction
regulation of RNA metabolic process
organelle fission
nucleic acid metabolic process
nuclear division
mitotic cell cycle
meiotic cell cycle
cytoskeleton organization
chromosome segregation
chromosome organization
cellular metabolic process
cellular aromatic compound metabolic process
Description
NES
p.adjust
C
Differentially expressed genes
WT cyd-1 RNAi Vs WT control RNAi
Downregulated
Upregulated
Unchanged
lim-7
log2(FoldChange)
-log10(padj)
D
Sheath cell structure scoring scheme (Phalloidin staining)
Normal
Mild defect
Severe defect
E
Sheath cell maker (lim-7p::gfp expression)
RNAi
Control
sys-1
daf-2
daf-16;daf-2
F
Normal
Reduced
Missing
lim-7p::gfp expression
.012
<.001
.006
RNAi
Control
sys-1
daf-2
daf-16;daf-2
G
Hermaphrodite spermatheca marker
(fkh-6::gfp expression)
RNAi
Control
cyd-1
daf-2
daf-16;daf-2
H
Normal
Reduced
Missing
fkh-6::gfp expression
<.001
<.001
RNAi
Control
cyd-1
daf-2
daf-16;daf-2
I
Sheath cell (Phalloidin staining)
RNAi
Control
cyd-1
WT Day-3
J
Normal
Mild defect
Severe defect
Sheath cell structure
RNAi
Control
cyd-1
WT (Day-3)

Figure S5

**Supplementary Figure 5.**

**(A-B)** Gene set enrichment analysis of differentially expressed genes in (A) *daf-2(e1370);rrf-3(pk1426);rde-1(ne219) V;unc-62p::rde-1(genomic)* or (B) *rrf-3(pk1426);rde-1(ne219) V;unc-62p::rde-1(genomic)* upon *cyd-1* KD as compared to control RNAi.

**(C)** Volcano plot showing the magnitude [$\log_2$(FC)] and significance [$-\log_{10}$(P value)] of the genes that are differentially expressed in L4 stage of *rrf-3(pk1426);rde-1(ne219) V;unc-62p::rde-1(genomic)* worms, grown on control or *cyd-1* RNAi.

**(D)** Sheath cell structure scoring scheme. Representative fluorescent images of phalloidin-stained (that marks the F-actin) gonads of worms (day-1 adult). The quality was categorized as normal, or mild or severe based on the structure and organization of actin filaments. Normal = No loosened or disorganized structure, mild = either loosened or disorganized actin filaments, severe= highly loosened or disorganized actin filaments. Yellow arrows point towards the defect.

**(E,F)** Representative fluorescent and DIC merged images of gonads showing *lim-7p::gfp* (that marks the sheath cells) expression in *daf-2(e1370)* and *daf-16(mgdf50);daf-2(e1370)* worms (day-1 adult) grown on control and *sys-1* RNAi. The gonadal arm is outlined for clarity. Scale bar 20 µm (E). Quantification of the normal, reduced or missing expression of *lim-7p::gfp* (F). Average of three biological replicates ($n \geq 18$ for each experiment). Two-way ANOVA-Tukey's multiple comparisons test.

**(G,H)** Representative fluorescent and DIC merged images of gonads showing *fkh-6::gfp* (that marks the hermaphrodite spermatheca) expression in *daf-2(e1370)* and *daf-16(mgdf50);daf-2(e1370)* worms (day-1 adult) grown on control and *cyd-1* RNAi. Spermatheca is outlined for clarity. Scale bar 20 µm (G). Quantification of the normal or missing expression of *fkh-6::gfp* (H). Average of three biological replicates ($n \geq 18$ for each experiment). Two-way ANOVA-Tukey's multiple comparisons test.

**(I,J)** Representative fluorescent images of phalloidin-stained (that marks the F-actin) gonads of WT worms (day-3 adult) grown on control and *cyd-1* RNAi. Scale bar 20 µm (I). Quantification of the normal or defective sheath cell structure (as per scoring scheme in Figure S5D) (J). Average of three biological replicates ($n \geq 13$ for each experiment). Unpaired *t*-test with Welch's correction.

Error bars are s.e.m. Experiments were performed at 20°C. Source data are provided in Table S1.

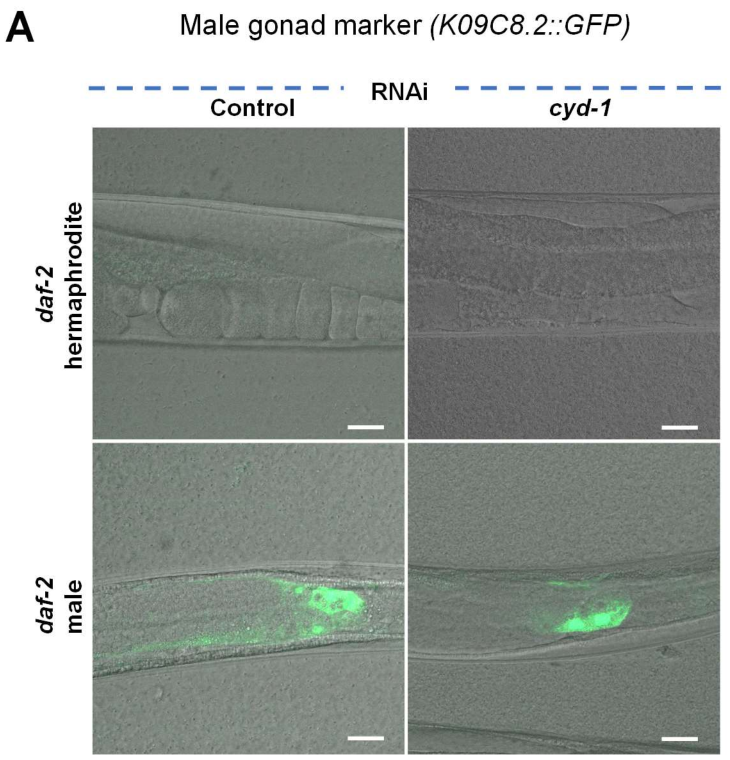


Figure S6

**Supplementary Figure 6.**

(A) Representative fluorescent and DIC merged images of gonads showing *K09C8.2::gfp* (male gonad marker) expression in *daf-2(e1370)* hermaphrodite or male worms (day-1 adult) grown on control and *cyd-1* RNAi. Scale bar 20 μm.
